# Dynamic temperature-sensitive A-to-I RNA editing in the brain of a heterothermic mammal during hibernation

**DOI:** 10.1101/288159

**Authors:** Kent A. Riemondy, Austin E. Gillen, Emily A. White, Lori K. Bogren, Jay R. Hesselberth, Sandra L. Martin

## Abstract

RNA editing diversifies genomically encoded information to expand the complexity of the transcriptome. In ectothermic organisms, including *Drosophila* and *Cephalopoda*, where body temperature mirrors ambient temperature, decreases in environmental temperature lead to increases in A-to-I RNA editing and cause amino acid recoding events that are thought to be adaptive responses to temperature fluctuations. In contrast, endothermic mammals, including humans and mice, typically maintain a constant body temperature despite environmental changes. Here, A-to-I editing primarily targets repeat elements, rarely results in the recoding of amino acids and plays a critical role in innate immune tolerance. Hibernating ground squirrels provide a unique opportunity to examine RNA editing in a heterothermic mammal whose body temperature varies over 30°C and can be maintained at 5°C for many days during torpor. We profiled the transcriptome in three brain regions at six physiological states to quantify RNA editing and determine whether cold-induced RNA editing modifies the transcriptome as a potential mechanism for neuroprotection at low temperature during hibernation. We identified 5,165 A-to-I editing sites in 1,205 genes with dynamically increased editing after prolonged cold exposure. The majority (99.6%) of the cold-increased editing sites are outside of previously annotated coding regions, 82.7% lie in SINE-derived repeats, and 12 sites are predicted to recode amino acids. Additionally, A-to-I editing frequencies increase with increasing cold-exposure demonstrating that ADAR remains active during torpor. Our findings suggest that dynamic A-to-I editing at low body temperature may provide a neuroprotective mechanism to limit aberrant dsRNA accumulation during torpor in the mammalian hibernator.

## INTRODUCTION

RNA editing promotes transcriptome diversity by expanding the coding capacity of the genome. The most common type of RNA editing is A-to-I editing, which is mediated by the Adenosine Deaminase that Acts on RNA (ADAR) family of enzymes. ADARs recognize dsRNA structures of at least 23 basepairs and deaminate up to 50% or more of their adenosines (Nishikura et al. 1991; Bass and Weintraub 1988). In mammals, there are two broad roles for mRNA editing: restricting the accumulation of dsRNA structures and altering codon sequences within coding regions. Mice deficient for *ADAR* die at embryonic day 13.5 due to systemic activation of the innate immune response (Mannion et al. 2014). Additionally in humans, mutations in *ADAR* result in the autoimmune disorder Aicardi-Goutieres syndrome, due to aberrant recognition of dsRNA and activation of innate immune signaling (Rice et al. 2012). In contrast, in mice loss of *ADARB1* results in seizures and lethality 2-3 weeks after birth due to loss of editing at a single recoding site in the glutamate receptor *GRIA2 (Brusa et al. 1995)*.

ADAR enzyme substrate specificity is sensitive to fluctuations in temperature because the stability of dsRNA structures are influenced by temperature (Wan et al. 2012). As temperatures decrease, short dsRNA helices are stabilized, resulting in an increased number of potential ADAR substrates. In *Drosophila* and *Cephlapod* species, reduced environmental temperatures leads to decreased body temperature and increased RNA editing (Buchumenski et al. 2017; Garrett and Rosenthal 2012; Savva et al. 2012; Yablonovitch et al. 2017b). In flies, decreased temperatures result in increased editing at a recoding editing site in the *ADAR* mRNA itself, as well as increased editing at 55 additional protein coding (CDS) sites (Savva et al. 2012; Buchumenski et al. 2017). The absence of RNA editing results in perturbed acclimation to temperature changes, suggesting an adaptive function for temperature-sensitive RNA editing (Buchumenski et al. 2017). Similarly, octopus species that reside in colder temperatures have higher rates of RNA editing of potassium channels, which is an adaptation that alters Kv1 channel kinetics to improve function in the cold (Garrett and Rosenthal 2012). While temperature-sensitive A-to-I RNA editing has been described in these ectothermic animals, it is unknown if such temperature-sensitive RNA editing occurs in mammals, which typically maintain a constant high body temperature (T_b_) despite environmental temperature fluctuations.

Hibernators provide a unique opportunity to assess whether temperature-dependent RNA-editing exists in mammals. Throughout winter, hibernating mammals cycle between extended periods of torpor at low T_b_ (>1 week, ∼4°C), and short (<12 hr) periods of arousal when T_b_ rapidly rises to the more typical mammalian temperature of ∼37°C (**Fig 1**), despite constant, near freezing environmental temperatures.

**Figure 1.**
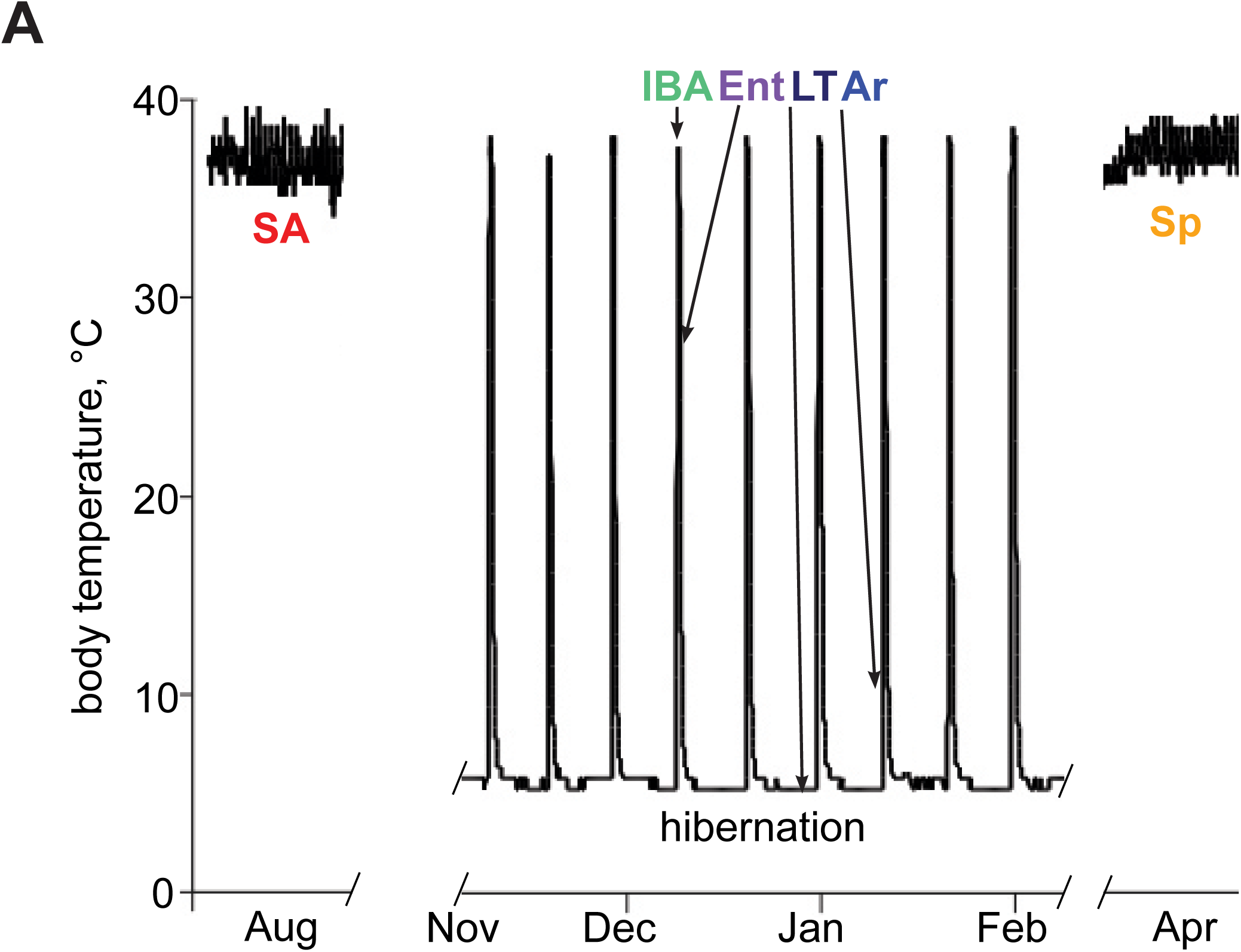
13-lined ground squirrels are heterothermic hibernators. (A) Schematic of sampling time-points selected for RNA-seq analysis. T_b_ for sampled animals: InterBout Arousal (IBA) 34.1°C +/- 2.9, Entrance (Ent) 25.4°C +/- 1.8, Late Torpor (LT) 5.9°C +/- 0.5, Arousing (Ar) 8.7°C +/- 2.1. T_b_ was assumed to be 37°C for Summer Active (SA) and SpD (Spring Dark) animals due to observed animal activity.

In this study we exploit the natural temperature fluctuations of hibernation to determine, if similar to ectotherms, RNA editing is increased by low temperature in these seasonal heterotherms. Neural tissue from three brain regions was selected for analysis because ADAR activity is typically found at high levels in the mammalian brain, ADAR recoding events occur in neural enriched transcripts including ion-channels (Tan et al. 2017; Behm and Öhman 2016; Yablonovitch et al. 2017a), and ion channels and other brain transcripts have been shown to be adaptively edited to alter function across a range of temperatures in ectotherms (Garrett and Rosenthal 2012; Yablonovitch et al. 2017b; Palladino et al. 2000). Our data demonstrate that ADAR-mediated A-to-I RNA editing occurs at thousands of sites in the brain transcriptome of hibernating ground squirrels while they are torpid at low T_b_.

## RESULTS

### Identification of RNA editing sites in squirrel brain tissues

Three brain regions, medulla, hypothalamus, and cerebrum, were selected for analysis of RNA editing due to their established roles in autonomic physiological processes, body temperature regulation and higher order cognitive processing, respectively. Furthermore, the medulla and hypothalamus remain relatively active compared to the cerebrum during the torpor phase of hibernation (Kilduff et al. 1990). A total of 90 paired-end, strand-specific RNA-seq libraries were generated from animals at six sampling points including two non-hibernating homeothermic stages: Summer Active (SA) and Spring Dark (SpD); and four of the heterothermic stages that characterize hibernation: Entrance (Ent), Late Torpor (LT), Arousing from torpor (Ar), and InterBout Arousal (IBA) (**Fig 1 and 2A**). These RNA-seq data were combined with data from testes and neonatal tissue samples to assemble a novel transcriptome that supplements the existing Ensembl annotations, which, because they are based largely on the automated recognition of protein sequence similarity, often lack complete mRNA annotations (e.g., missing 5’ and 3’ UTRs; see Materials and Methods). Over 86.1% +/- 2.6 of the RNA-seq reads recovered from the brain samples were uniquely alignable, with 85.4% +/- 1.2 of those uniquely mapped reads overlapping exon annotations defined in the novel transcriptome in contrast to only 53.4% +/- 2.5 mapping to exons when annotated with the existing Ensembl transcriptome (**S1 Fig**).

To identify and characterize RNA editing sites, single nucleotide variants were found using the Genome Analysis ToolKit (GATK) best practices for calling variants from RNA-seq (DePristo et al. 2011) (**Fig 2**, **S2 Fig**). Variants were called individually for each of the 90 samples, merged, filtered based on quality metrics, and assigned to a strand based on the strandedness of the paired end alignment orientation (**Fig 2, S2A Fig**). In total, A-to-G was the most common variant detected, accounting for 26.3% of the single nucleotide polymorphisms (SNPs) identified. But a substantial proportion identified were non-A-to-G variants (**S2B Fig**). Because the animals used for this study were obtained from largely outbred populations they are expected to harbor genetic variation; the excess of transition substitutions (C-T, G-A, T-C, A-G) over transversions is consistent with this expected genetic diversity (**S2B Fig**). Despite the large number of polymorphisms recovered, however, we observed a 2.2 fold increase of A-to-G variants over the expected distribution derived from T-to-C variants, an enrichment not observed for other mismatch pairs (**S2C Fig**) and consistent with ADAR-mediated RNA editing activity.

**Figure 2.**
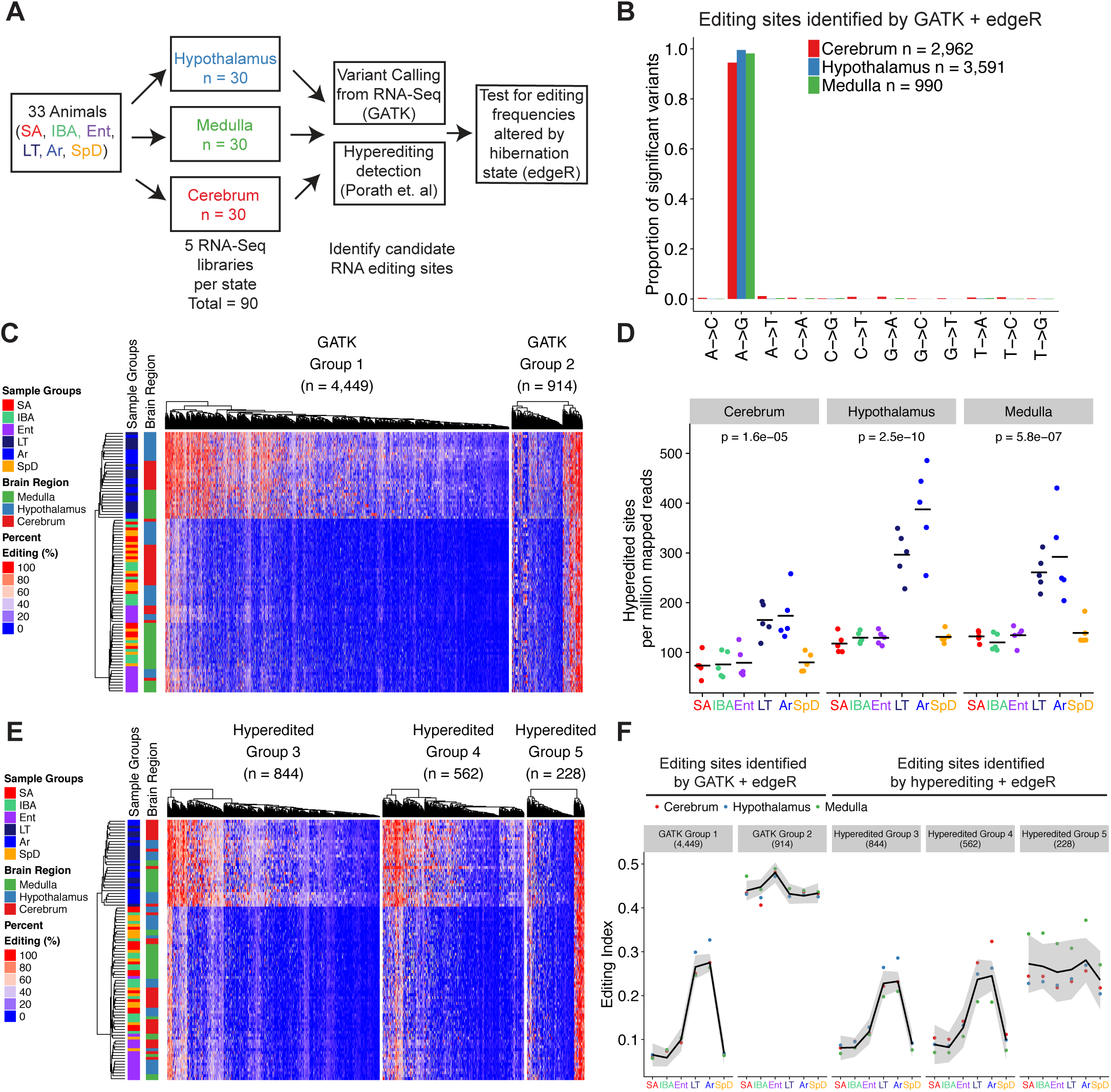
A-to-I RNA editing is widespread and increases at low temperature during torpor and arousing from torpor. **A**. RNA-seq library information and RNA editing site identification approach, see Fig 1 for full description physiological stages. **B.** Proportion of variants identified by the GATK pipeline for each possible mismatch type that have significantly changed editing frequencies across hibernation sampling timepoints determined via GLM analysis (FDR < 0.01). **C.** Heatmap depicting editing frequencies defined as the percentage of G containing reads over the reads with either A or G. Group 1 or Group 2 sites were defined by k-means clustering (k = 2 classes), and within each Group, editing sites and individual samples were ordered by hierarchical clustering. **D.** The number of editing sites detected by the hyper-editing approach in each brain region (panel) and hibernation state (color) normalized to the total number of uniquely mapped reads. P-values were determined via ANOVA. **E.** Heatmap of editing frequencies for sites identified by the hyper-editing approach that are significantly changing across hibernation states. Groups were defined with k-means clustering (k = 3 classes) and sites were ordered with hierarchical clustering. **F.** Editing index for significant sites identified by the GATK or hyper-editing approach across sampled hibernation states. The editing index is defined as the sum of all G containing reads for all significant sites over the sum of all reads at significant sites. Lines are fit using a loess method and shaded area indicates standard error estimates derived from the t-distribution.

There are 59 A-to-I editing sites that are conserved across mammals (Pinto et al. 2014). We recovered 51 of these 59 A-to-I RNA editing sites in the set of A-to-G variants identified prior to filtering to exclude sites with extremely high or low variant allele frequencies (**S1 Table**). These sites included amino acid recoding sites in the glutamate receptor *GRIA2* (Gln560Arg in ground squirrel, *ENSSTOG00000024582*) and the serotonin receptor, *HTR2C* (Ile156Val, Ile156Met, Asn158Ser, Ile160Val, in ground squirrel, *ENSSTOG00000028990*) (**S3 Fig**). In addition, we also recovered several mouse (*n* = 100) and human (*n* = 212) editing sites previously identified in a curated database of RNA editing sites (Ramaswami and Li 2014) (**S1 Table**). Together, these observations demonstrate that, in addition to detecting polymorphic sites in the ground squirrel brain transcriptome, we also recover previously characterized RNA editing events in the set of A-to-G variants.

The 13-lined ground squirrel genome currently lacks high quality transcriptome and SNP annotations, complicating accurate assignment of variants as *bona fide* RNA-editing events using RNA-seq data. Commonly used methods for excluding polymorphisms from candidate RNA editing sites rely on SNP databases and therefore cannot be applied to remove SNPs from our candidate RNA-editing sites (Ramaswami et al. 2013). However, we reasoned that SNPs should be randomly distributed throughout our study population and would therefore not be enriched within samples derived from a specific hibernation state. We enumerated the number of reads with variant and reference alleles for each site and applied a statistical model to identify variants whose allele frequencies are significantly associated with a hibernation state. We applied a general linear model based ANOVA-like test (edgeR) to each set of the 12 possible mismatch types (i.e., A-to-G, A-to-C, T-to-G, etc) and assessed changes in variant allele frequencies while accounting for changes in the coverage at the site. From the original set of A-to-G variants (*n* = 179,295), we identified subsets of variants that are significantly associated with hibernation state in the medulla (*n* = 990), hypothalamus (*n* = 3,591), and cerebrum (*n* = 2,962) respectively (FDR < 0.01). Moreover, these variants accounted for greater than 94% of the significant sites identified in each of these regions (**Fig 2B**). The remaining minority population of variants (G-to-A, C-to-T, T-to-C, and A-to-T) were primarily transition mismatch variants that likely represent polymorphisms that passed our statistical cut offs due to the large number of variants tested for significance. SNPs are distributed equally across both DNA strands, in contrast to RNA editing sites, which are specifically localized to the transcribed strand. The lack of enrichment for T-to-C variants in the set of hibernation state-dependent variants demonstrates that the significant A-to-G variant sites are not likely to be SNPs, but rather result from A-to-I RNA editing events mediated by the ADAR family of deaminases.

### Widespread increases in A-to-I RNA editing during torpor

The editing frequencies of the sites that differed significantly by hibernation stage were clustered and visualized to ascertain their pattern of editing. Two clear patterns were evident after sites were classified into two groups by k-means clustering (**Fig 2C**). Group 1 comprises 82.9% of the editing sites and displays increased editing frequencies during late torpor and arousing from torpor where T_b_ at collection was 5.9°C +/- 0.5 and 8.7°C +/- 2.1, respectively (**Fig 2F**). Samples from animals in late torpor were at depressed T_b_ (<30°C) for >85% of the length of the previous torpor bout (7.7 +/- 2.2d) compared to samples from arousing hibernators which were at low T_b_ for the entire torpor bout (10.2 +/- 2.0d), and still had low T_b_ (<9°C) but were beginning to rewarm. The highest editing index is reached as the animals begin to arouse from torpor, indicating that editing frequencies are increased by both low temperature and the amount of time spent with T_b_ at low temperature.

In contrast to Group 1 sites, Group 2 sites were not enriched for any particular hibernation stage, suggesting that sites in this group are either editing sites that have variable editing frequencies independent of hibernation physiology or are contaminating SNPs that were not eliminated by our statistical approach. We also clustered the variant allele frequencies for the small number (n = 127) of significant non-A-to-G variants and found that these variants were not enriched for any particular hibernation stage, consistent with these minor sites likely being SNPs (**S4 Fig).**

### Hyper-editing events are enriched in Late Torpor and Arousing stages

To corroborate the observed increases in RNA-editing during torpor we employed a second approach to identify RNA editing sites. ADAR enzymes can edit adenosines in a dsRNA helix to a degree that generates hyper-edited RNAs that are unable to be aligned to the genome after sequencing (Carmi et al. 2011). To identify these hyper-editing events, we used a previously described approach that can reliably recover A-to-I editing with a low rate of false-positive SNP identification (**S5 Fig**) (Porath et al. 2014). This approach identifies hyper-edited sites by converting all As in each unaligned read and the genome to G, prior to alignment. For successfully aligned reads, the original genome and read sequence are recovered and alignments with a large number of mismatches are retained as hyper-edited (see Materials and Methods). This approach has also been used to identify RNA editing events in non-model organisms that lack curated databases of genetic variation (Liscovitch-Brauer et al. 2017; Porath et al. 2017b).

To assess the specificity of this pipeline for recovering A-to-I editing events, we independently identified hyper-edited sites for all possible mismatch types. The majority (93.7%) of hyper-edited sites identified were A-to-G mismatches (**S6A Fig**), with the remaining sites comprising T-to-C and other transitions, with very few transversions. Thousands of unique A-to-G hyper-editing sites were identified (*n* = 85,737); the number of these sites increased strongly during torpor and as animals aroused from torpor in all three brain regions, consistent with the increased editing frequency identified using the GATK + edgeR approach (**Figs 2D and S6B Fig**). In contrast to the low T_b_ LT and Ar stages, the number of hyper-editing sites did not fluctuate across the homeothermic stages (SpD, SA) or warm heterothermic stages (IBA, Ent) (Figs. 2D and S6C and S6D Fig).

We next computed the editing frequency for the hyper-edited sites using only reads aligned to the unmodified genome (i.e. without changing A-to-G) and tested these sites for significant changes across hibernation states, in the same manner as the GATK + edgeR approach. We recovered 1,634 significant hyper-editing sites (FDR < 0.01) (**Fig 2E**) and used k-means clustering to define three groups of hyper-edited sites. Sites in Groups 3 and 4 displayed similar increased editing frequencies when animals were in late torpor or arousing from torpor, whereas Group 5 sites were largely invariant across hibernation states, similar to the Group 2 sites identified by the GATK + edgeR pipeline (**Fig 2F**).

The Group 3 and 4 editing sites identified by the hyper-editing approach and the Group 1 sites identified by the GATK pipeline were designated as cold-enriched editing sites (**Fig 3A**). The sites identified by both the GATK and the hyper-editing pipeline that were not called significant in the ANOVA-like edgeR analysis were designated as a set of constitutively edited sites with editing frequency independent of temperature or hibernation physiology (**Fig 3B and 3C**). We required these non-significant constitutively edited sites to be identified by both pipelines to reduce the likelihood of misclassifying a SNP as an editing site.

**Figure 3.**
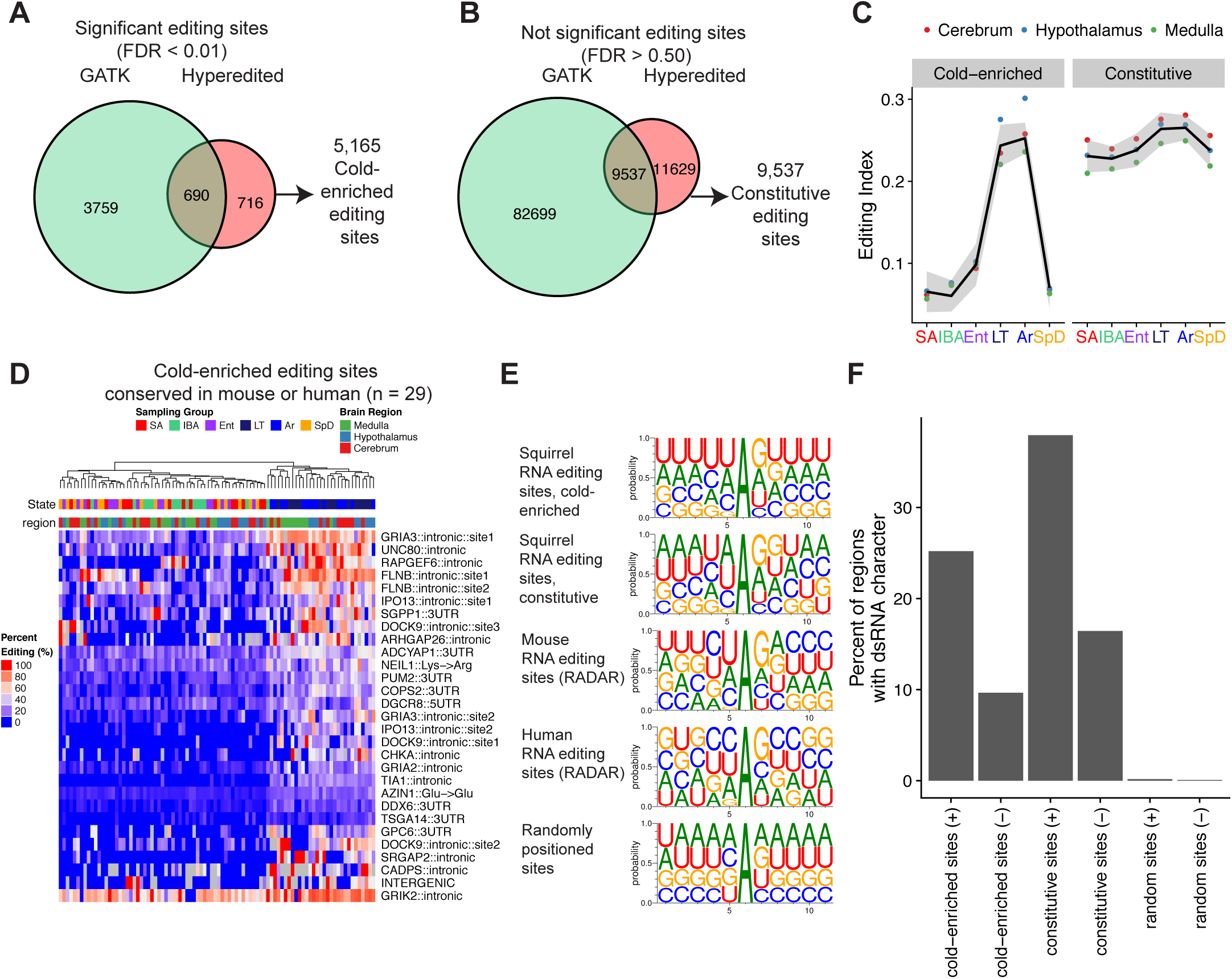
Novel editing sites have an ADAR sequence motif and are enriched in regions with dsRNA character. **A.** Euler diagram depicting the number of significant editing sites detected by the GATK or hyper-editing pipeline. Cold-enriched sites were defined as the union of the two sets of significant sites. **B.** Euler diagram illustrating the set of constitutively edited sites defined as the non-significant editing sites (FDR > 0.5) identified by both the GATK and the hyper-editing pipeline. Only sites identified by both methods were selected to reduce the likelihood of misclassifying a SNP as an editing site. **C.** Editing index for cold-enriched and constitutively edited sites. **D.** Heatmap depicting editing sites conserved in either mouse or human, ordered by hierarchical clustering. Predicted locations of these sites are based on annotations in the RADAR database. **E.** Sequence logo of sequences surrounding all cold-enriched or constitutive editing sites (+/- 5 nucleotides) or editing sites identified in mouse in the RADAR database. A set of randomly positioned sites was also generated by randomly selecting an A nucleotide position from transcripts containing editing sites. **F.** Quantification of dsRNA character of each editing site via *blastn* by aligning +/- 100 nucleotide regions surrounding editing sites, to the reverse complement of a +/- 2 kb region surrounding the editing site or to the end of the transcript. Sites with an e-value < 0.1 with alignment over the editing site are considered paired dsRNA. As a control, the reverse complement, denoted as (-), of each +/- 100 nt region were also quantified and the number of second best alignments are shown.

### Conservation of mRNA editing sites

We compared the set of common significantly edited sites (*n* = 5,165) identified by the hyper-editing or the GATK pipeline (**Fig 3A**) to previously characterized mouse or human editing sites collected in the RADAR database of editing sites (Ramaswami and Li 2014). 27 cold-enriched sites were previously identified in humans, and 4 identified in mouse, with 2 of the 31 sites being identified in both the mouse and human sets. This set of 29 sites includes intronic editing sites in the glutamate receptors; *GRIA2, GRIA3,* and *GRIK2,* UTR editing sites in the RNA binding proteins *PUM2* and *DGCR8,* and the recoding site in the DNA glycosylase *NEIL1* (Lys242Arg) (**S1 Table**) (Yeo et al. 2010). The editing frequencies for these 29 conserved sites were clustered, and additionally demonstrated the same pattern of enriched editing frequencies in late torpor and arousing animals as observed when clustering all of the significant editing sites (**Fig 3D**).

### Sequence and structural features of cold-enriched mRNA editing sites

We next examined the sequence and structural context of the cold-enriched sites and the set of editing sites whose editing frequencies did not vary across samples. Both cold-enriched and constitutively edited sites have a sequence preference for depletion of guanosine 5’ of the edited adenosine and enrichment for guanosine 3’ of the site, a pattern not observed for randomly selected adenosines (**Fig 3E**). This sequence preference is similar to the sequence context preferred for ADAR enzymes for editing and to the sequence preferences for previously identified RNA editing sites in mouse and humans in the RADAR database (Eggington et al. 2011).

ADAR deaminates adenosines in dsRNA structures of at least 23 basepairs in length that form by pairing with editing complementary sequences (ECS) (Nishikura et al. 1991). Identifying ECS sequences is challenging as they can be located proximal to the editing site, in distal regions such as an intronic ECS that basepairs with an exon, or mediated through intermolecular interactions (Reenan 2005; Bass and Weintraub 1988; Ramaswami et al. 2015). To identify potential ECS sequences in our ground squirrel data, a 201 nt region surrounding each editing site was aligned to a larger region (4001 nt) surrounding the editing site. Alignments matching the reverse complement of the larger region represent sequences capable of basepairing with the editing site and were enumerated for cold-enriched, constitutive, and a set of shuffled control sites (**Fig 3F**). A large percentage of cold-enriched (25.2%) and constitutive (38.0.%) editing sites had identifiable ECS sequences, in contrast to reverse complemented negative controls (9.6% and 16.4% for cold-enriched and constitutive respectively) or shuffled control sites (0.2% and 0.1%). Taken together these results demonstrate that the cold-enriched and constitutive editing sites both have sequence and structural features consistent with known A-to-I editing sites.

### mRNA editing events are enriched in SINE element repeats

A-to-I editing events in mammals are predominantly located in SINE-element derived tandem inverted repeats (Porath et al. 2017a). A much smaller proportion are located in exonic sequences, with very few cases of editing sites that recode amino-acids, in contrast to *Drosophila* or *Cephalopods*, where recoding events are more common (Liscovitch-Brauer et al. 2017). We therefore next assessed the genomic distribution of the ground squirrel brain editing sites. The cold-enriched editing sites are primarily localized in retained intron sequences, similar to the distribution of RNA editing sites in human brain (**Fig 4A**) (Hwang et al. 2016). Constitutively edited sites have a similar distribution to the cold-enriched sites, indicating that the temperature-sensitive RNA-editing sites are not preferentially localized to exonic or coding sequences (**Fig 4B**). 90.1% of the constitutive and 82.5% of the cold-enriched editing sites reside in SINE repeat regions, with the majority being STRID repeat elements (**Figs. 4C and 4D**). In the squirrel lineage, the STRID SINE repeats are derived from tRNA, similar to ID-elements in mice and analogous to 7SL-RNA derived Alu elements in primates (Vassetzky and Kramerov 2013; Churakov et al. 2010). These results demonstrate that the temperature-sensitive editing sites are not preferentially localized to coding or exonic regions, and are primarily repeat-derived.

**Figure 4.**
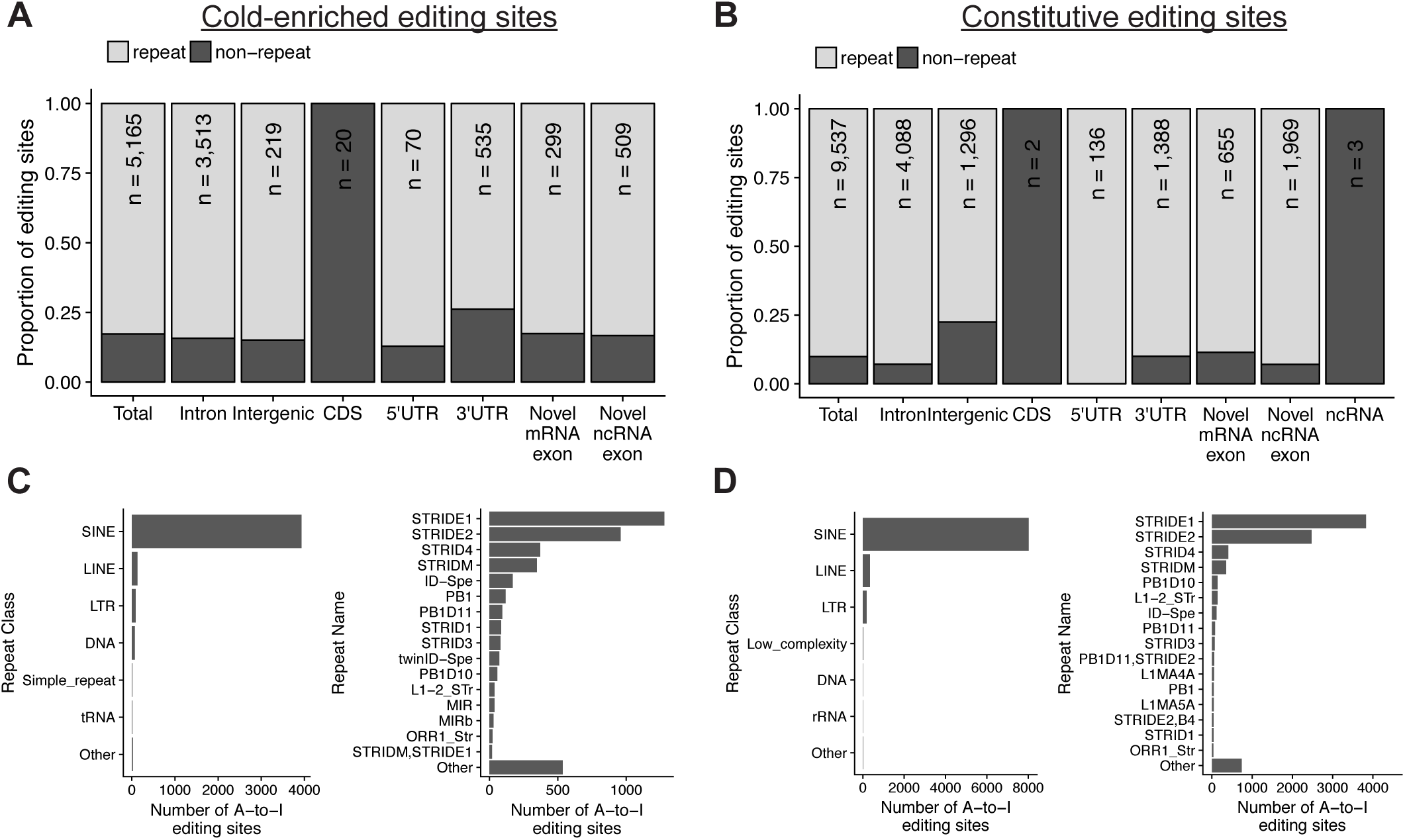
Editing sites are predominantly located in SINE elements in intronic sequences. **A.** Genomic distribution of cold-enriched editing sites. Editing sites within exons from novel transcripts that fall within protein coding genes but do not overlap existing CDS exons are annotated as novel mRNA exons, otherwise they are annotated as intronic. Editing sites within novel annotations that do not overlap any known protein-coding annotations are classified as overlapping novel ncRNA exons **B.** Distribution of constitutively edited sites. **C.** Number of cold-enriched editing sites within repeat classes and repeat families defined by repeatMasker annotations. **D.** Repeat classification for constitutively edited sites.

### Functional Impact of Editing

We next assessed the potential functional impacts of the cold-enriched editing on specific mRNAs. We used snpEFF to annotate the predicted effects of A-to-G substitution based on gene annotations from Ensembl. The majority of sites reside in either intron or intergenic regions, and are not predicted to overlap with splice acceptor or donor sequences (**Fig 5A**). Just 13 sites are predicted to be either moderately or highly deleterious, 12 of these are recoding events and one is in a splice acceptor site (**Fig 5B and S2 Table**). Additionally, 8 CDS editing sites are silent, and do not recode an amino acid.

**Figure 5.**
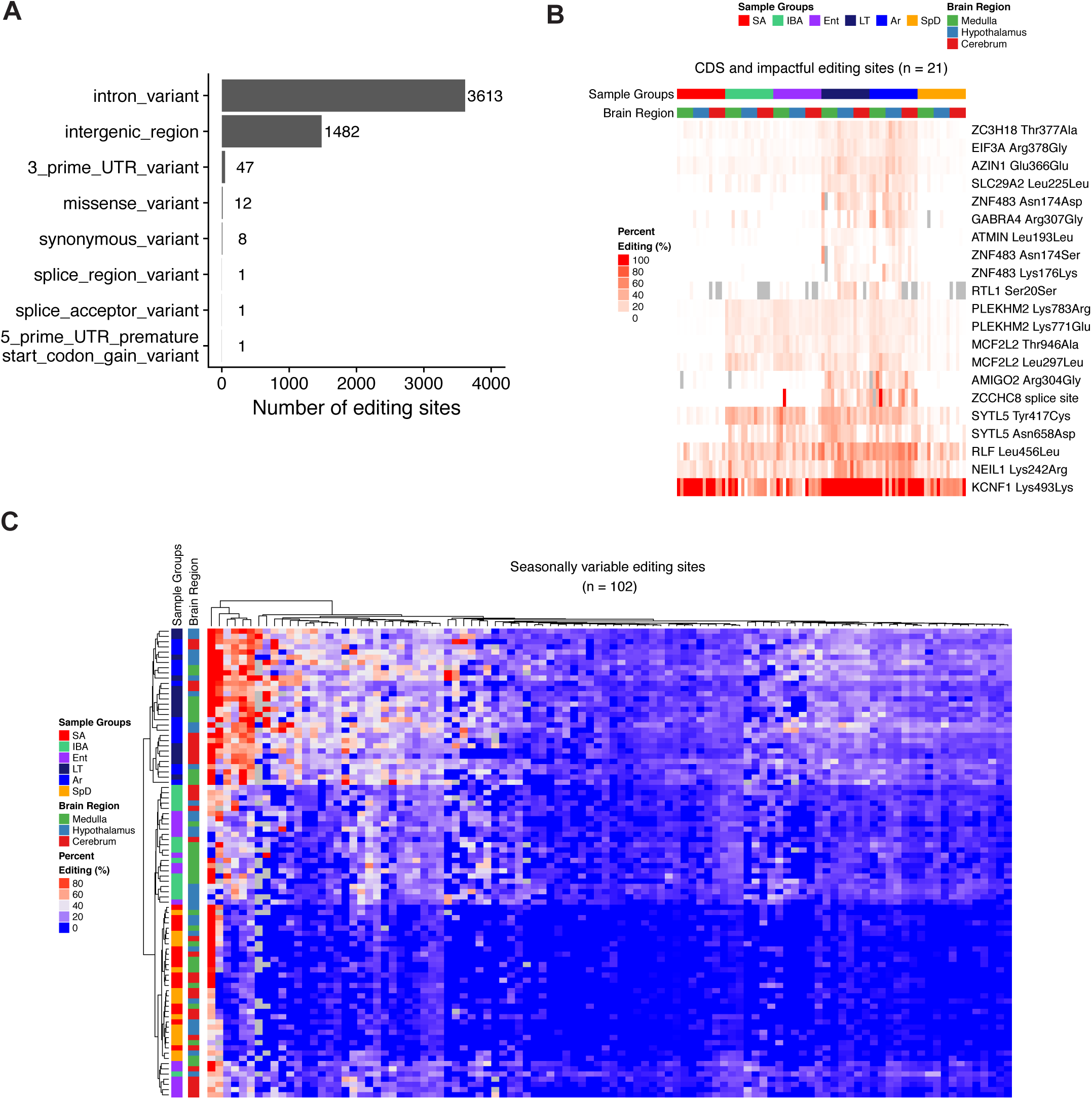
Predicted functional consequences of editing. **A.** Summary of predicted impacts of cold-enriched editing events as defined by SnpEff using Ensembl annotations. **B.** Heatmap depicting RNA editing frequencies for editing sites in CDS regions or predicted to be deleterious by SNPeff analysis (Moderate or High impact). Sites without read coverage are colored grey. Columns are ordered by sample group and rows are ordered by hierarchical clustering. **C.** Heatmap depicting cold-enriched sites with significant seasonally altered editing frequencies (FDR < 0.01). Comparisons were made between heterothermic warm samples (Ent and IBA) and homeothermic warm samples (SA and SpD). Rows and columns are clustered by hierarchical clustering.

Examining the editing frequencies for the predicted deleterious sites revealed that the recoding sites in *PLEKHM2*, *SYTL5* and *MCF2L2* were additionally edited during IBA and Ent stages, suggesting that the editing of these sites is seasonally regulated (**Fig 5B**). We therefore examined the cold-enriched editing sites to additionally classify sites with seasonally altered editing frequencies. Seasonally regulated editing sites were identified by comparing the warm heterothermic stages IBA (34.1°C +/- 2.9) and Ent (25.4°C +/- 1.8) to warm homeothermic stages SpD (37°C) and SA (37°C). From this analysis we identified 102 sites with significantly increased editing frequencies in IBA and Ent compared to SpD and SA (FDR < 0.01) (**Fig 5C**), demonstrating that a small proportion of the cold-enriched sites (1.9%) are also edited in a seasonally dependent manner.

We selected 10 of the putative RNA-editing sites for validation in a subset of the ground squirrels by dideoxy sequencing of the corresponding liver genomic gDNA (**S7-9 Figs.**). The selected sites included nine of the temperature-sensitive sites, seven predicted to be deleterious and two intronic sites, and the universally edited Gln560Arg site in *GRIA2*. Additionally a Group 2 site identified by the GATK pipeline that was predicted to be a SNP rather than an editing site was selected as a positive control for detecting homozygous vs. heterozygous SNPs via this method. cDNA and gDNA sequencing demonstrated that none of the predicted RNA editing events were actually SNPs, and additionally validated the predicted SNP in *DKK3* mRNA (**S9 Fig**). Validated recoding events include the splice-acceptor site alteration in *ZCCHC8 (ENSSTOG00000001683),* a component of the Nuclear EXosome Targeting complex (NEXT) and a recoding event in *Z3CH18* (Thr377Ala, *ENSSTOG00000022431*) a scaffolding protein that links the NEXT complex to the Cap Binding Complex to promote exonucleolytic cleavage of snRNA and replication dependent histone mRNAs (Meola et al. 2016). We also verified the recoding site in *EIF3A* (Arg378Gly, *ENSSTOG00000002512*), the largest subunit of EIF3 which is required for EIF3 assembly and translation initiation (Wagner et al. 2014). In addition, editing caused the amino acid replacement at the previously described Lys242Arg site in *NEIL1 (ENSSTOG00000005781),* a DNA glycosylase whose substrate specificity is altered by the arginine substitution (Yeo et al. 2010), and a recoding event in one subunit of gamma-aminobutyric acid receptor, *GABRA4* (Arg307Gly, *ENSSTOG00000001951*), which is a ligand-gated chloride channel for the main inhibitory neurotransmitter in mammalian brains.

The splice-acceptor editing site in the *ZCCHC8* mRNA occurs in a retained intron. We therefore investigated if cold-enriched RNA editing at this site modulated the frequency of intron retention. The editing frequency at the splice-acceptor site increases up to 21.0% in late torpor and 53.3% during arousal compared to a near absence of editing observed in the other states (**S10 Fig)**. However, the relative proportion of the retained intron did not increase in the cold animals (LT and Ar), suggesting that editing of this mRNA occurs post-transcriptionally and therefore does not impact its splicing.

### mRNA editing frequencies increase during torpor

Our observation that editing frequencies increased in the LT and Ar animals compared to the IBA and Ent animals could be the result of either RNA editing during entrance into torpor as T_b_ falls below 23°C, or of editing throughout the 1-2 week torpor period while T_b_ is maintained at 5-6°C. To distinguish between these possibilities, cerebrum samples were taken at an earlier stage of torpor, when T_b_ is less than 30°C for 1.2 +/- 0.3d. Editing frequencies for intronic sites in *FBXW7* and *CDH9* were determined by dideoxy DNA sequencing of cDNA from five early torpor animals for comparison to five LT animals. These sites were chosen because they had nearly undetectable editing in the IBA and Ent animals, but had greatly increased A-to-G substitutions during LT and Ar, providing the greatest dynamic range with which to assess the change in editing frequency across the torpor bout (**Fig 6A**). Because the editing frequencies in early torpor were higher than those observed at Ent, some editing has already occurred in the first 1-2 days with T_b_ < 23°C, but the editing frequencies at both sites were significantly lower during early torpor than late torpor, indicating that editing occurred across the torpor bout despite the continuously low T_b_. The increased editing frequency during torpor and arousal from torpor is unlikely to be explained by changes in ADAR abundance, as we observe only an 19.5 +/- 6.3% increase in ADAR mRNA in LT compared to IBA (**S11 Fig)**. This small increase is most likely explained by a relatively increased stability of ADAR mRNA compared to the mRNA pool (Grabek et al. 2015), and is unlikely to affect the ADAR protein pool because initiation of translation, and thus translation of any newly edited mRNAs, is blocked at the low T_b_ of torpor (van Breukelen and Martin 2001). Increased editing frequencies across the torpor bout could in principle be the result of either de novo editing by ADARs, or alternatively, selective stabilization of mRNAs edited prior to ET. To distinguish between these possibilities we examined the normalized read counts containing unedited (A) or edited (G) nucleotides across hibernation stages. During LT and Ar the normalized abundance of edited transcripts increased while the abundance of unedited transcripts decreased proportionally (**Figs. 6B and S12 Fig**). Given the absence of transcription at the low T_b_ of torpor (van Breukelen and Martin 2002), these findings favor a model in which *de novo* ADAR activity throughout the period of low T_b_ results in increasing editing frequencies.

**Figure 6.**
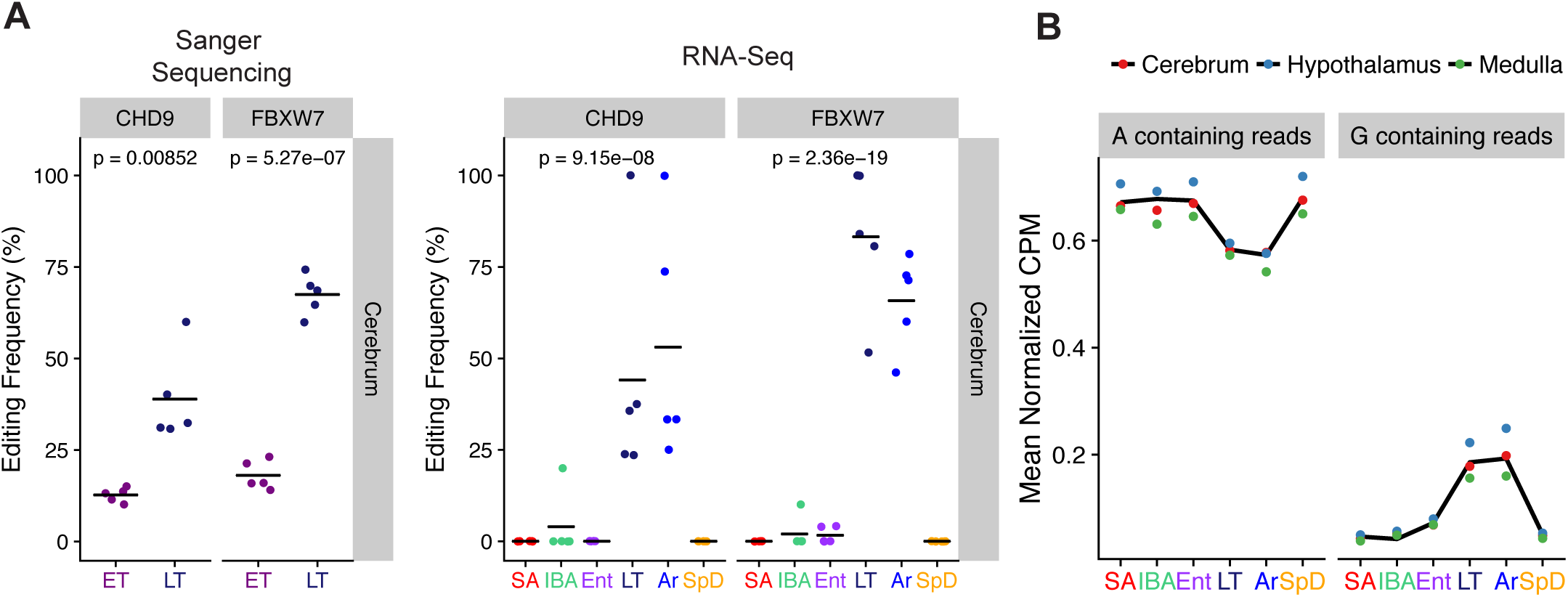
A-to-I editing frequencies increase during torpor. **A.** Editing frequencies for intronic sites in *FBXW7* and *CHD9* determined by sanger sequencing from cerebrum samples (n = 5) at early torpor (ET) or late torpor (LT). Editing frequencies determined by RNA-seq are also shown for comparison. **B.** For all cold-enriched sites the average normalized abundance (counts per million) for reads containing reference (A) or edited (G) nucleotides is shown.

## DISCUSSION

Our results demonstrate that widespread RNA editing occurs in three brain regions of a mammalian hibernator during torpor, when body temperature remains near freezing for multiple days. Temperature-sensitive RNA editing has been observed previously in *Drosophila* species exposed to acute temperature changes (Buchumenski et al. 2017; Yablonovitch et al. 2017b), and in Cephalopods, where ocean temperatures are negatively correlated with editing frequencies in a potassium channel mRNA across multiple species (Garrett and Rosenthal 2012). In both cases, increased A-to-I editing contributes to temperature adaptation, and is associated with increased editing in coding regions. In this study we find that ADAR-mediated RNA editing is also greatly increased in hibernating ground squirrels during torpor, when the animal’s temperature hovers near freezing for several days. The degree of editing at most of these sites increases with increasing time spent cold and then returns to baseline within 3hr of temperature restoration to 37°C during spontaneous arousal (**Figs. 2 and 6**), although some (1.9%) edited sites persist throughout the torpor-arousal cycle (**Figs. 5B and 5C)**. Only 20 of the hibernation edited sites with either pattern lie within coding regions, and just 12 of these would recode the corresponding protein. Instead, the vast majority of the edited sites reside in SINE-family interspersed repeats, as is typical of non-hibernating, homeothermic mammals (Porath et al. 2017a).

It is clear from our data that the dominant effect of the greatly enhanced RNA editing during the cold phase of hibernation is to target sequences that are likely to form dsRNA rather than to diversify the proteome for cold adaptation. Nevertheless, it remains formally plausible that one or more of the small set (*n* = 12) of edited sites predicted to recode amino acids (**Fig 5B**) could improve the corresponding protein’s function in the cold, as documented for the K_v_1 potassium channel in the octopus (Garrett and Rosenthal 2012), and thus be adaptive for hibernation. Interestingly, five of the 12 recoding events occur in edited sites that appear to be enhanced seasonally rather than strictly by temperature (**Fig 5B**), specifically at two sites in *PLEKHM2* and *SYTL5*, and one site in *MCF2L2*. These may be particularly important to help neurons function in the cold, because the level of editing remains consistently elevated throughout the torpor-arousal cycle, including during interbout arousals when edited transcripts can be actively translated into protein (Frerichs et al. 1998). Conversely, the seven remaining potential recoding events are less likely to substantially affect their corresponding protein pools because translational initiation is arrested at temperatures below 18°C (van Breukelen and Martin 2001) and the proportion of transcript that is edited quickly declines as T_b_ recovers to 37°C during IBA (**Figs. 2 and 6**), when translation resumes and the bulk of protein synthesis during hibernation occurs. For these transcripts, recoded proteins synthesized from cold-edited mRNAs are expected to reach their greatest concentrations just as euthermic T_b_ is restored rather than during the time of greatest need for their cold-adapted function, i.e., as T_b_ declines during entrance into torpor and throughout multiple days at low T_b_ in torpor. These kinetics taken together with the low editing frequencies for recoded sites (<40%) suggest that only a small fraction of the proteins made from the cold edited transcripts would be recoded.

Because transcription is largely suppressed during the near-freezing temperatures of torpor (van Breukelen and Martin 2002), ADAR enzymes must be active for edited sites to accumulate (**Fig 6**). ADAR activity during torpor provides a mechanism to destabilize dsRNA structures that form at lower temperatures, and by restricting the accumulation of dsRNA, RNA editing could prevent inappropriate activation of innate immune sensors upon rewarming as the animals arouse from torpor (Liddicoat et al. 2015). ADAR activity during torpor could also promote the nuclear retention of subsets of transcripts in via *P54NRB,* which recognizes inosine containing RNAs and sequesters them within the nucleus (Zhang and Carmichael 2001). The majority of cold-enriched editing that we observed in the hibernators occurs in transcripts with retained introns (**Fig 4A**), which are unlikely to produce a functional protein product. Nuclear retention of such transcripts could be used to prevent wasteful translation of inappropriately processed mRNAs generated during temperature transitions in the torpor-arousal cycle.

Hibernators are unique among mammals in their ability to allow body temperature to fall to near freezing and remain there for days to weeks at a time with no evidence of irreversible cellular damage or loss of organismal function on arousal (Dave et al. 2012). Given that homeothermy is an evolutionarily recent invention found in birds and mammals, the ability to maintain cellular function and integrity while cold in hibernation is likely a retained ancestral trait (Lovegrove et al. 2014). To date there is largely evidence against (Villanueva-Cañas et al. 2014) but little evidence for (Matos-Cruz et al. 2017) genetically encoded cold-adaptation of proteins that could support function at low temperature during hibernation. mRNA editing has the potential to cause adaptive changes in proteins that leave no signature in the genome; this mechanism is used for temperature adaptation of proteins in ectotherms (Garrett and Rosenthal 2012; Savva et al. 2012; Buchumenski et al. 2017). Here we demonstrate for the first time rampant, temperature-dependent RNA editing during hibernation, with most sites falling outside of protein coding regions. While we cannot rule out that the few protein recoding events observed in this study are adaptive for improved function in the cold, the bulk of the editing, as typical of homeothermic mammals studied previously, is directed towards interspersed repeats that engage in dsRNA formation. Blocking and marking regions of dsRNA at low temperature represses activation of the innate immune response (O’Connell et al. 2015) which is known to occur (Bouma et al. 2010), and also likely to be adaptive, during hibernation.

## MATERIALS AND METHODS

### Tissue Collection

Tissues were collected at precise timepoints based on T_b_ (**Fig 1**) from five ground squirrels in each of six distinct physiological stages. Animals from two homeothermic and four heterothermic (hibernation) phases were used (**Fig 1**). Under deep isoflurane anesthesia, animals were euthanized via exsanguination and perfused with ice cold saline. The brain was extracted from the skull, and then the brainstem and cerebellum were removed. The medulla and hypothalamus were dissected; for this study, these two regions and the remaining telencephalon plus diencephalon (excluding the hypothalamus, henceforth referred to as cerebrum) were snap frozen in liquid nitrogen and stored at -80°C until RNA extraction.

### RNA-seq

Either the entire frozen tissue (medulla, hypothalamus), or 100 mg of tissue pulverized under liquid nitrogen with a mortar and pestle (cerebrum) were homogenized in ice cold TRIzol Reagent (Invitrogen) on ice using a polytron (Brinkman), and then purified using the Direct-zol RNA MiniPrep Kit (Zymo Research), following the manufacturer’s recommendations. RNA was assessed for quantity (NanoDrop, Thermo Scientific) and quality (RIN > 8; Bioanalyzer, Agilent Technologies) and then 1 µg was submitted to the University of Colorado Denver Genomics and Microarray Core for library preparation and sequencing. Strand-specific RNA-seq libraries were constructed using the TruSeq Stranded mRNA Library Prep Kit (Illumina); paired-end sequence reads were collected for each sample after multiplexing via Illumina HiSeq 2500 (cerebrum, 10 samples/lane) or 4000 (hypothalamus and medulla, 15 samples/lane). Samples (replicates 1-5 from each physiological state) were assigned to a sequencing lane by multiplexing all of the first replicates from each state, followed by the 2nd replicates etc., to avoid batch effects and assure that roughly equivalent numbers of individual libraries representing each state were spread among the sequencing lanes.

### Transcriptome Assembly

A custom transcriptome assembly was built using all 90 brain, plus three neonatal and two testes libraries. The neonatal libraries were constructed from three regions of a P1 neonate, head, thorax and abdomen. The testes libraries were prepared from two aliquots of RNA isolated from one adult male testes during spring recrudescence following hibernation. The 95 RNA-seq libraries were aligned first to the 13-lined ground squirrel mitochondrial DNA sequence (Hampton et al. 2011) and then the remaining reads were aligned to SpeTri2.0 (Ensembl release 88, retaining contigs >=10000nt) with HISAT2 (v 2.0.4, (Kim et al. 2015)). BAM files were filtered to remove reads mapping to more than one genomic location (bamtools, v 2.4.0) and duplicates (Picard Tools, v 1.83), and the remaining reads were used for guided transcriptome reconstruction using StringTie (v. 1.3.3b, with options --rf, -j 3, -c 3) (Pertea et al. 2015; Kim et al. 2015). Transcriptomes were then merged to generate a single assembly using TACO (v. 0.7.2, with option --filter-min-expr st to 5.0) (Niknafs et al. 2016). The reconstructed transcriptome was used to define splice junctions for STAR mapping and annotating editing sites.

### RNA editing detection with the GATK pipeline

Candidate RNA editing sites were identified by following the GATK pipeline for variant calling from RNA-seq data as described in https://gatkforums.broadinstitute.org/gatk/discussion/4067/best-practices-for-variant-discovery-in-rnaseq. RNA-seq libraries were trimmed with cutadapt (v.1.8.3) (Martin 2011) (cerebrum) or Illuminas bcl2fastq (medulla and hypothalamus) to remove Illumina truseq adapters, then aligned to the SpeTri2.0 (Ensembl 85) genome assembly supplemented with the transcript assembly annotations built as described above. Alignment was performed using STAR (v.2.5.1b with default parameters) in two-pass mode as recommended by GATK (Dobin et al. 2013). Duplicate alignments were marked with MarkDuplicates from Picard (v.2.7.0), and read alignments over splice junctions were split into independent alignments using SplitNTrim from GATK (v3.5-0-ge91472d). Variants were called using HaplotypeCaller (-stand_call_conf 20.0, -stand_emit_conf 20.0), filtered to remove sites with Qual by Depth (QD) < 2.0 or Fisher Strand Score (FS) > 30. Variants were then merged and used as input for base recalibration. Variant calling was then rerun with the updated base recalibration table. This process was repeated twice to establish proper base recalibration as recommended by GATK for samples without a known set of SNPs. Final variant calls were then merged and filtered to remove sites with Depth (DP) < 20. The strandedness of the variant was determined by calculating the mean strand bias (positive or negative stranded reads / total number of reads) determined based on the read alignment strand and mate (R1 or R2), and averaged over all the libraries. Variants with > 0.80 positive or negative strand bias were assigned to a strand, whereas others were excluded as ambiguous. Reference and variant alleles were counted at each site using only reads with unique alignments, MAPQ > 10, base quality scores >= 20, not marked as duplicate, secondary, QC failed, or mate-non-mapped alignments. Additionally reads with variant sites located in the first 6 nucleotides were also excluded to avoid biases from random hexamer priming. Variants with extreme variant allele frequencies were then filtered to remove any variants that were not present at least >5% variant allele or <95% frequencies in any of the 90 libraries to enrich for variants with dynamic allele frequencies across samples.

### Hyper-editing detection

Hyper-edited regions were identified following the methods described in a published method and were implemented with custom Python scripts (Porath et al. 2014). Briefly, reads that did not align after two-pass STAR mapping were subjected to filtering to exclude low-quality reads and reads with highly repetitive sequences, as previously described. Following filtering, A nucleotides were converted to Gs in forward stranded reads (from paired mate R2) and mapped to a genome also with As changed to Gs to capture sense alignments, or a genome with Ts changed to Cs to capture antisense alignments. Similarly in reads from paired mate R1 (reverse stranded) Ts were changed to Cs and aligned to genomes with either T to C or A to G changes. Alignments were performed using BWA (v0.7.10) with no gaps and only two allowed mismatches (Li and Durbin 2009). Successful alignments were then processed to identify mismatches between the original read and genome sequences. Alignments with high quality mismatches passing a stringent set of filters to reduce misalignment artifacts were retained as hyper-edited reads as previously described (Porath et al. 2014)]. The described procedure was repeated for each of the 12 possible mismatch types to determine the specificity of this approach for A-to-G mismatches. Hyper-edited regions were defined in each read as the region spanning the first and last mismatch. Hyper-edited clusters were defined by merging all overlapping regions or those located within 20 nucleotides. Clusters and sites supported by fewer than two reads were discarded. Editing frequencies were computed for hyper-edited sites by using reads aligned during STAR two-pass mapping to the unmodified genome with reference and variant alleles counted as described for the GATK pipeline.

### Differential editing analysis

Reference allele counts at each candidate editing sites were normalized using library sizes derived from the total number of exonic alignments in each library and scaled with TMM normalization using the R package, edgeR (v3.18.1). Normalization factors were then propagated to the variant allele counts. A GLM model was next constructed, ∼0 + Animal + Hibernation Stage:Allele, whereby Animal = animal id, Hibernation Stage = IBA, Ent, Lt, Ar, Spd, or SA, and Allele = A or G. P-values were obtained using glmLRT, with contrasts set to test for sites with variable G counts across sampled hibernation stages while controlling for changes in A counts within each animal (McCarthy et al. 2012). Sites with FDR < 0.01 were considered significant. This procedure was independently applied for all possible pairwise allelic combinations (i.e. A-to-T, A-to-G, A-to-C, T-to-A, T-to-G, etc). Editing frequencies were visualized using the R package ComplexHeatmap (v1.14.0) (Gu et al. 2016). K-means clustering was performed on editing frequencies mean-centered and scaled across all samples using the flexclust package (1.3-4) with the kmeans++ initialization function (Leisch 2006). Sites without sufficient read coverage to compute editing frequencies were set to zero for k-means classification. Individual samples and editing site ordering in the displayed heatmaps were determined by hierarchical clustering of euclidean distances with the complete linkage method.

### Snpeff

The potential functional impacts of the cold-enriched editing sites were annotated with SnpEff (v.4.3b) using Ensembl 85 annotations (Cingolani et al. 2012).

### RNA editing site validation

PCR primers (**Table S3**) were designed to amplify gDNA or cDNA surrounding candidate editing sites. Six cold-enriched editing sites were validated by both gDNA and cDNA sequencing of cold (Ar) and warm (SpD) animal samples from either liver for gDNA or brain tissue for cDNA, (*ZCCHC8*, *EIF3A, ZC3H18, AMIGO2, FBXW7, CHD9)*, in addition to the positive control site in *GRIA2* **(S9 Fig**). The negative control *DKK3* site (**S9 Fig**) and three cold-enriched editing sites were validated by gDNA sequencing from liver gDNAs alone (*GABRA4*, *NEIL1*, and *ZNF483*) (Data not shown). PCR products were subjected to dideoxy sequencing. Peak heights from chromatograms were determined with the ThermoFisher QC app tool (https://apps.thermofisher.com/apps/dashboard/#/).

### Differential gene expression and splicing analysis

Read counts were computed using featureCounts from the subread package (v1.4.4) using the custom transcriptome annotations (Liao et al. 2014). Lowly expressed genes were filtered if there were not at 2 counts per million in at least 4 samples. Normalized FPKM values were calculated using edgeR (rpkm()). Differentially expressed genes were identified using an ANOVA-like test for any variation across hibernation state. Relative exon usage values were computed using DEXSeq (v.1.16.10).

### Editing site conservation

Genome coordinates for human (hg19) and mouse (mm9) editing sites were downloaded from the RADAR database (Ramaswami and Li 2014). These coordinates were converted to mm10 coordinates using UCSC liftover chain files and the liftOver tool. Squirrel RNA editing sites were converted to mm10 coordinates using the UCSC liftover chain and liftOver tool, and shared editing sites were identified using the R package valr (v0.3.1) (Riemondy et al. 2017). Editing sites conserved in mammals were taken from the supplementary data of (Pinto et al. 2014) and compared to the squirrel editing sites as described above.

### Sequence Motifs

Sequences 5 nucleotides upstream and downstream of each editing site were extracted and sequence logos were computed with weblogo (v3.5.0) (Crooks et al. 2004). A set of negative control shuffled editing sites were constructed for the cold-enriched sites by randomly selecting an adenosine nucleotide in the same transcript as each editing site. Random sites were drawn from exonic sequences if the editing site was exonic, otherwise the sites were drawn from pre-mRNA transcript coordinates.

### RNA structure predictions

Editing sites were classified as falling within dsRNA using an approach as previously described (Li et al. 2009). Briefly, a 201 nucleotide region centered on the editing site was aligned to the reverse complement of a larger surrounding region (4001nt) to identify sequence capable of basepairing to the editing site. blastn (v2.2.29, -strand “plus” -word_size 7 -evalue “0.1”) was used in the two sequence alignment mode. Alignments were filtered to keep alignments with e-value < 0.01 and alignment regions >20 nucleotides overlapping the editing site. Negative controls were generated by aligning the region surrounding the editing site to the same stranded larger region and selecting the second best alignment using the aforementioned criteria.

### Software and Data availability

The RNA-seq raw data, transcriptome assembly, and processed data will be made available through GEO. The RNA editing pipelines were implemented as a snakemake pipeline (Köster and Rahmann 2012) and custom scripts for hyper-editing detection were written in Python and C++. Statistical procedures, data-processing, and visualizations were implemented in R. The pipeline, scripts, and a link to a UCSC genome browser trackhub with the RNA-seq data can be found at https://github.com/rnabioco/rnaedits.

## ACKNOWLEDGEMENTS

We acknowledge support from the RNA Bioscience Initiative at the University of Colorado School of Medicine (K.A.R., A.E.G., E.W., J.R.H. and S.L.M.) and the National Institutes of Health (R21 NS088315 to S.L.M. and J.R.H). We thank K. Diener in the University of Colorado Cancer Center Genomics Resource for expert assistance with DNA library construction and sequencing (supported by NIH grant P30 CA046934) and E. Epperson for brain dissection (supported by NIH grant R01 HL089049 to S.L.M).

## Supplemental Figure Legends

**S1 Fig. Quality control of RNA-seq libraries.** (A) Total number and percentage (B) of uniquely alignable reads from each RNA-seq library. (C) Percent of reads overlapping exons, introns, or unannotated per library, when defined by existing Ensembl annotations or the transcriptome built in this study (stringtie + taco).

**S2 Fig. Variant identification workflow and distribution of detected variants.** (A) Schematic for variant calling based on GATK best practices for variant calling from RNA-seq. (B) Distribution of variants detected by GATK pipeline. (C) Fold enrichment of variants detected over variants detected for each complement mismatch type.

**S3 Fig. Editing Frequencies for conserved HTR2C and GRIA2 editing sites.** (A) Editing frequencies for HTR2C and GRIA2 CDS recoding sites conserved in mammals.

**S4 Fig. Significant non-A-to-G variants are not enriched for late torpor or arousing states.** (A) Heatmaps of variant allele frequencies for significant non-A-to-G mismatches. Note that the rows displayed in each heatmap represent different variant sites. Rows and columns are ordered by hierarchical clustering.

**S5 Fig. Pipeline for identifying hyperedited regions from unaligned reads** (A) Hyperediting pipeline implemented as described by Porath H. et al. 2014. Mean +/- sd of reads, alignments, or sites of A-to-G conversions are shown.

**S6 Fig. Hyperedited sites are primarily A-to-G variant type and enriched in Arousing and Late Torpor** (A) Number of unique hyperedited sites detected for each possible allele type following approach described in Porath et al. Only sites supported by at least 2 reads are shown. (B) Distribution of shared unique hyperediting sites identified in each sampling group. (C) Unique hyperedited clusters identified in each brain region for each variant type. Clusters located within 20 nucleotides were merged. (D) Unique hyperedited sites identified in each brain region for each variant type.

**S7 Fig. Heatmap of RNA editing frequencies for editing sites selected for gDNA and cDNA validation.** (A) Predicted effects of the editing events are annotated beneath each gene name. The *GRIA2* R/G editing site was selected as a known editing site as a positive control. The *DKK3* site is a group 2 site identified by the GATK pipeline that was predicted to likely be a SNP and therefore selected as a control to confirm the sanger sequencing approach. Sites without read coverage are colored grey.

**S8 Fig. Abundance of edited transcripts selected for validation.** (A-J) Normalized FPKM values for genes containing editing sites selected for validation by sanger sequencing. FDR values are derived from an ANOVA-like test for differential expression across sampled hibernation states.

**S9 Fig. Dideoxy sequencing validation of editing sites.** Chromatograms show the edited nt (arrow) +/- 3 neighboring nt for each site from cerebrum cDNA and liver genomic DNA from one Ar (cold hibernator) and one SpD (warm euthermic) individual ground squirrel. Six of the cold-edited sites recovered in our analysis are shown in addition to the fully ADAR edited (+) control GRIA2 site. The bottom right *DKK3* (-) control shows liver genomic DNA sequence recovered from four individuals, two homozygous and two heterozygous, at the GATK group II site that was predicted to be a SNP.

**S10 Fig. RNA editing does not disrupt *ZCCHC8* splicing at exons 11 and 12.** (A) Editing frequency of an editing site that is predicted to disrupt a splice-acceptor site (B) UCSC browser snapshot of RNA-seq coverage across retained intron that contains the editing site. Coverage tracks are taken from a single library from each brain region. Ensembl transcript annotations are displayed. (C) Relative usage of intron containing predicted splice-acceptor disrupting editing site or flanking exons. p-values were computed as an ANOVA-like test for relative usage changes correlated with any sample group using DEXSeq.

**S11 Fig. mRNA abundance of ADAR family members across hibernation stages.** (A) Normalized FPKM for *ADAR*, *ADARB1*, and *ADARB2*. FDR values derived from ANOVA-like test for differential expression across sampled hibernation states.

**S12 Fig. Normalized counts for reads containing G increase during late torpor and arousing.** (A) Heatmap of normalized read counts (CPM) for reads containing A (reference) or G (edited) nucleotides. Columns for G containing reads are ordered by hierarchical clustering. Columns for A containing reads are ordered based on the column orders for samples in the G containing read heatmap for ease of visualization.

S1 Table. Editing sites conserved across mammals, mice, or humans.

S2 Table. Summaries for constitutive and cold-enriched editing sites.

S3 Table. Primer sequences for editing site validation by cDNA and gDNA Dideoxy sequencing.

