## Supplementary Materials for "Dynamic temperature-sensitive A-to-I RNA editing in the brain of a heterothermic mammal during hibernation"

**Supplemental Figure 1**

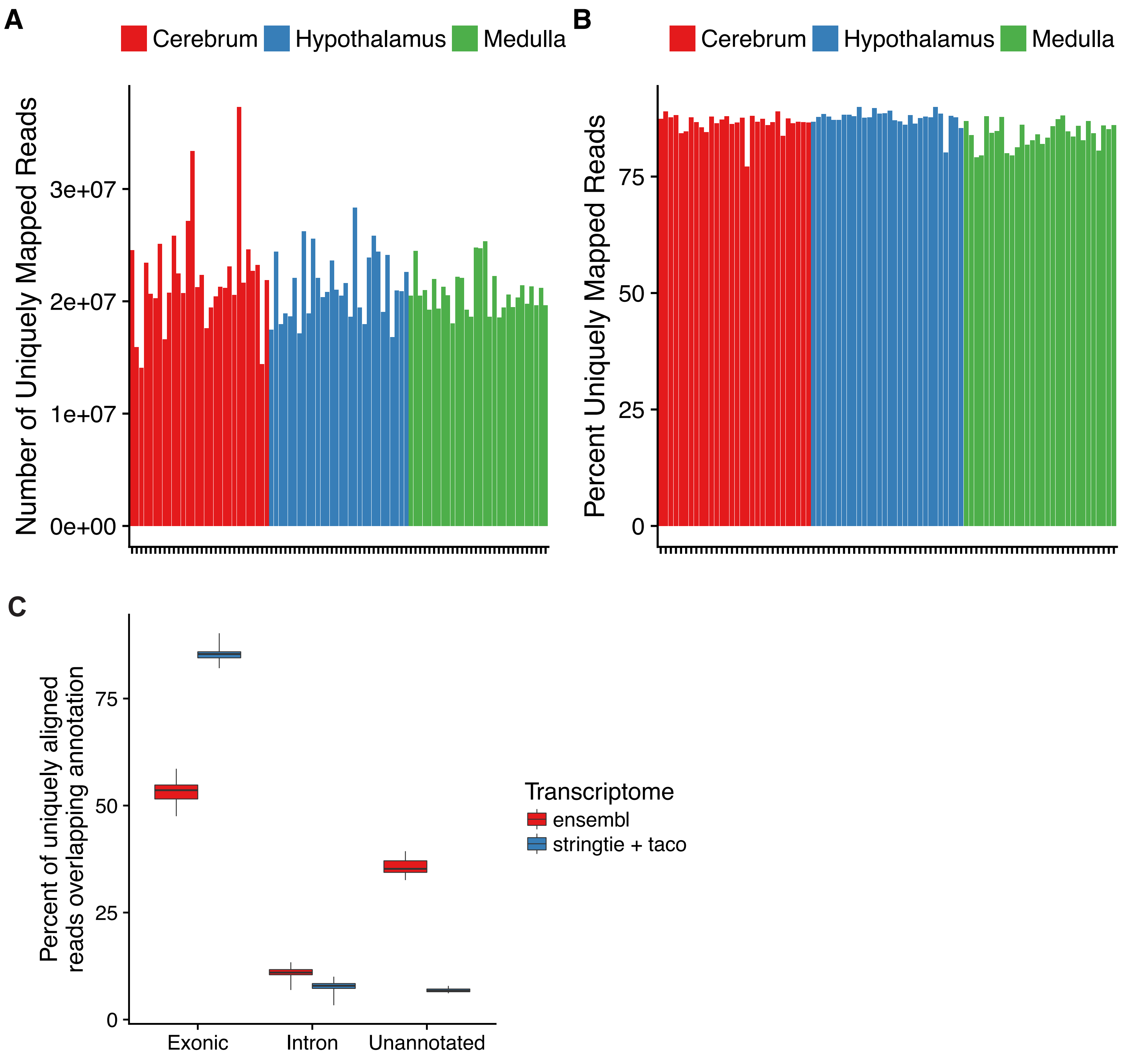

**Supplemental Figure 2**

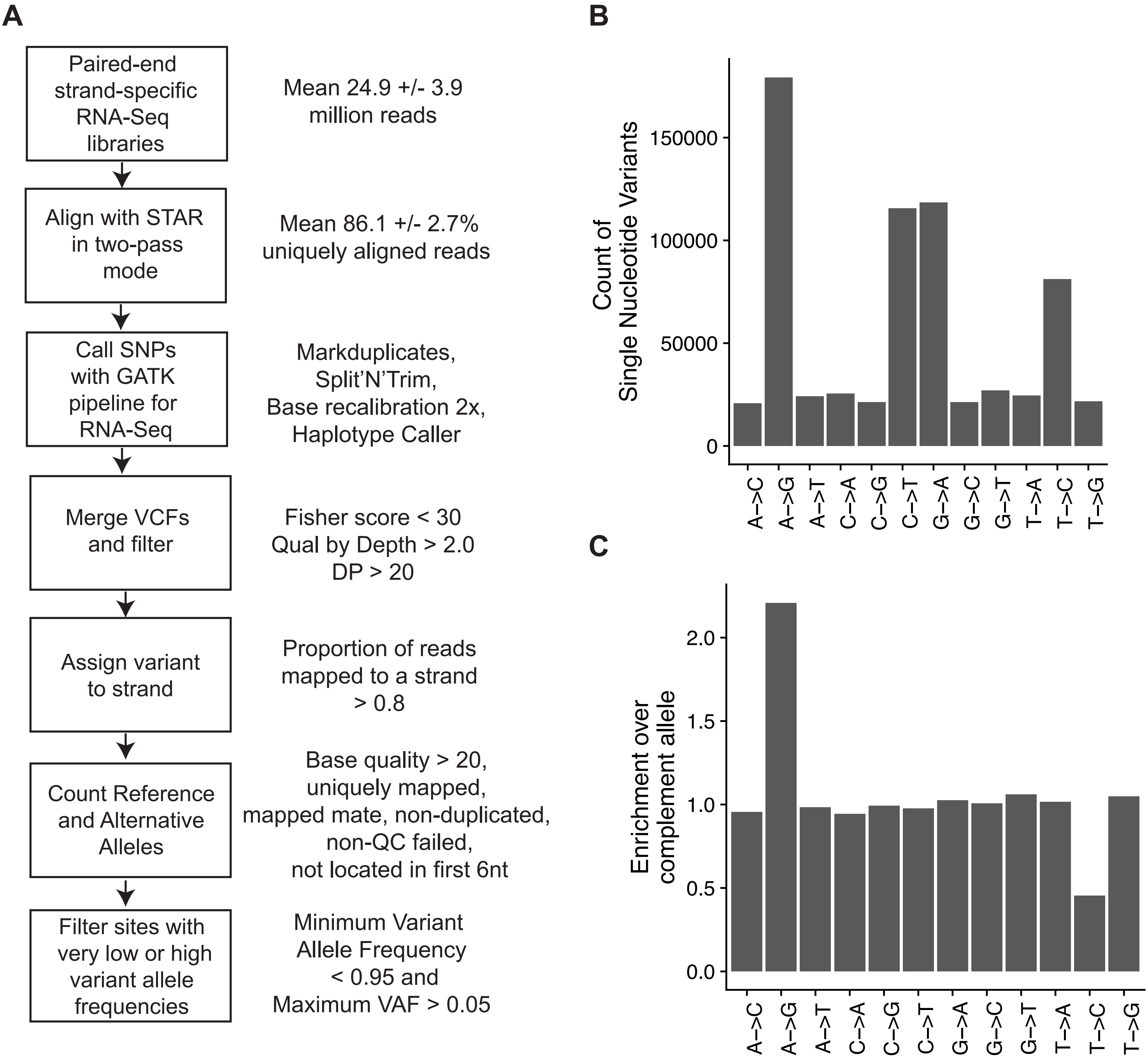

**Supplemental Figure 3**

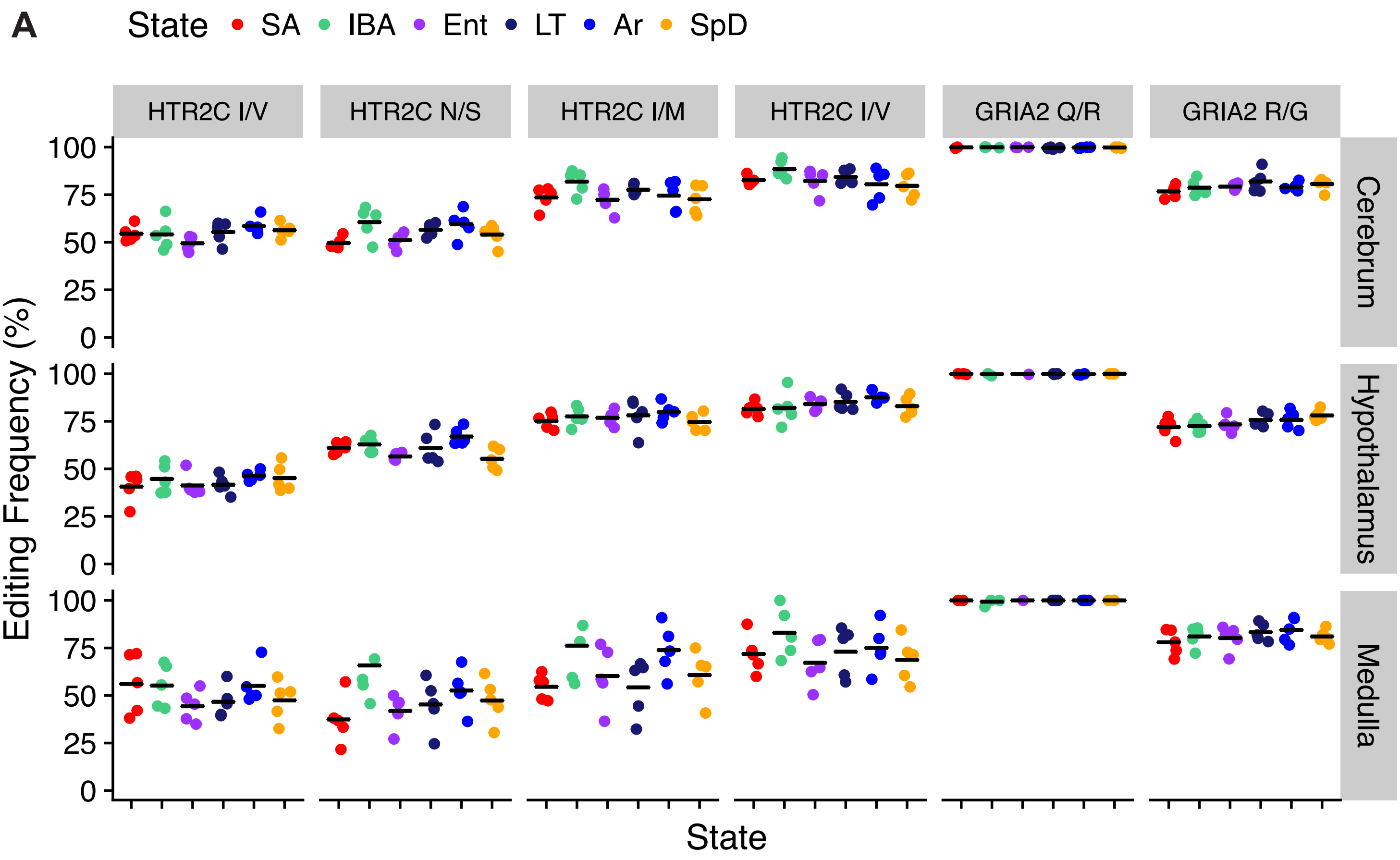

### Supplemental Figure 4

## A

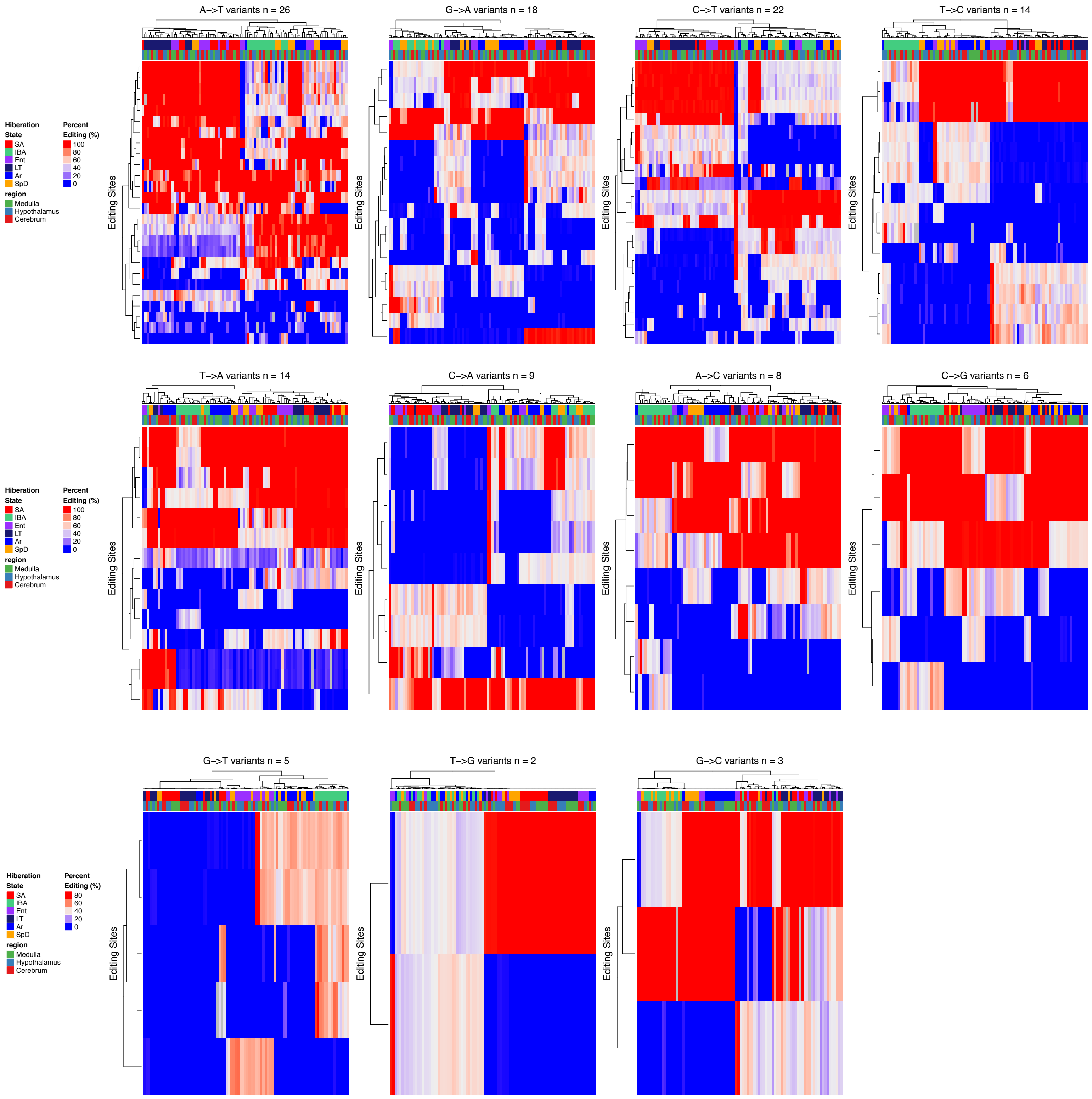

**Supplemental Figure 5**

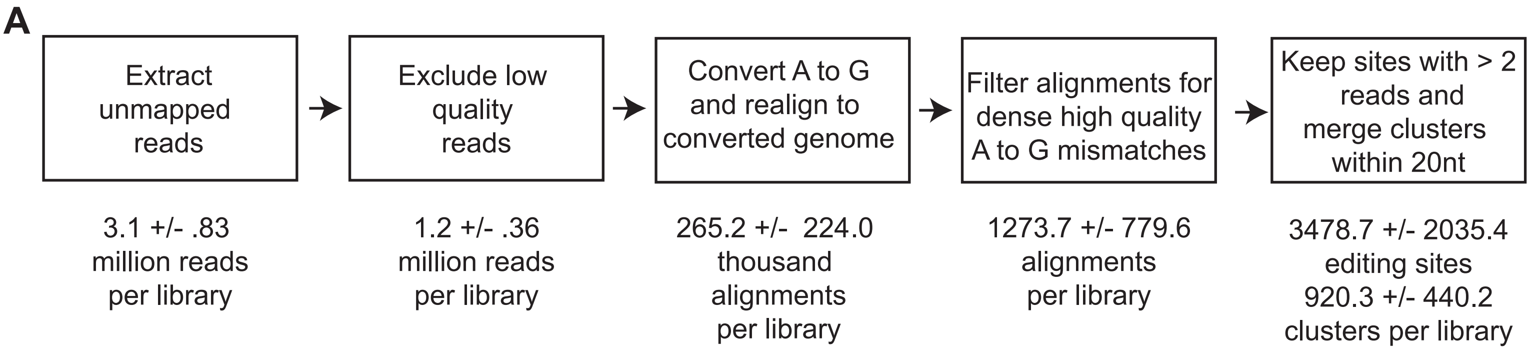

**Supplemental Figure 6**

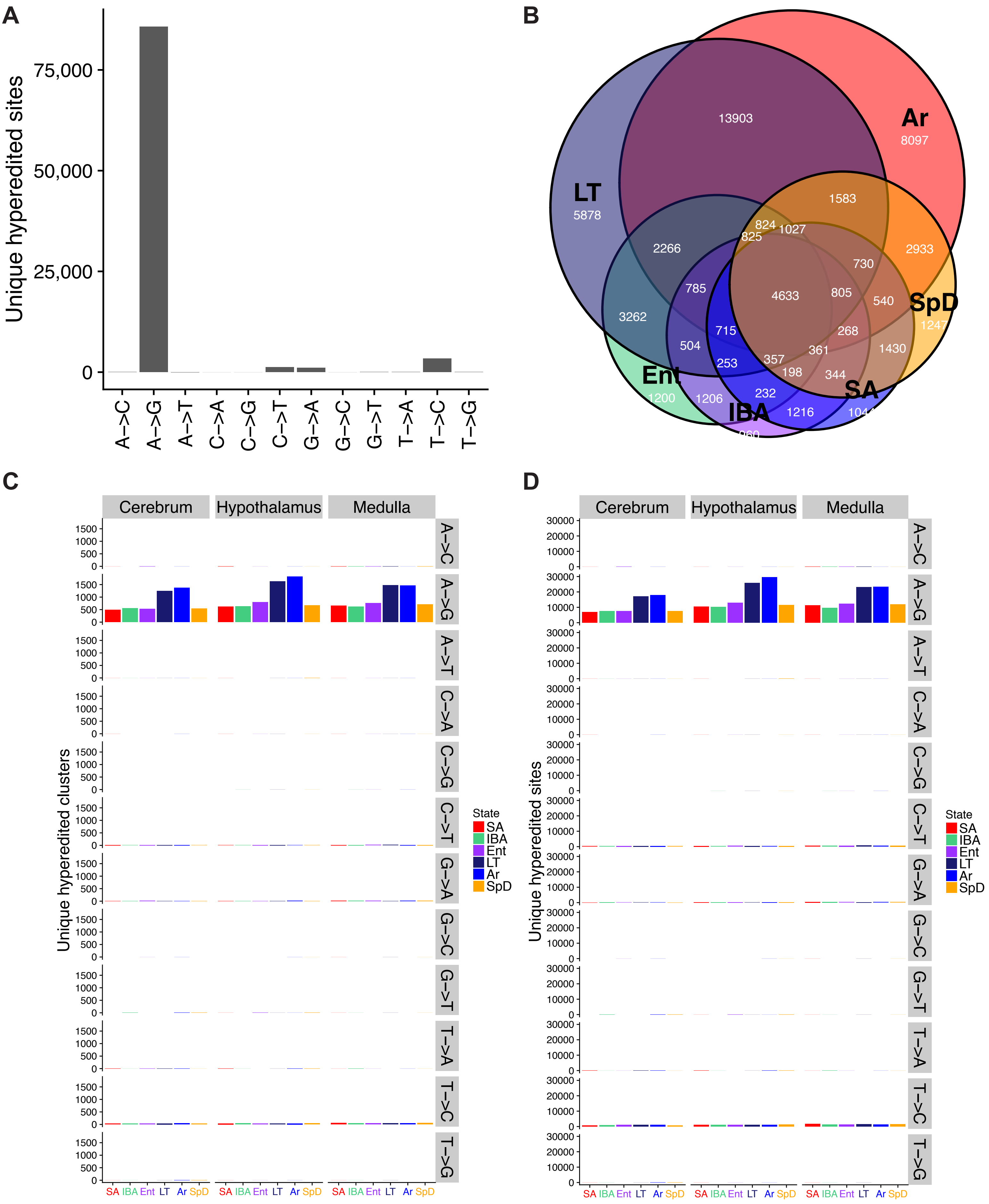

#### Supplemental Figure 7

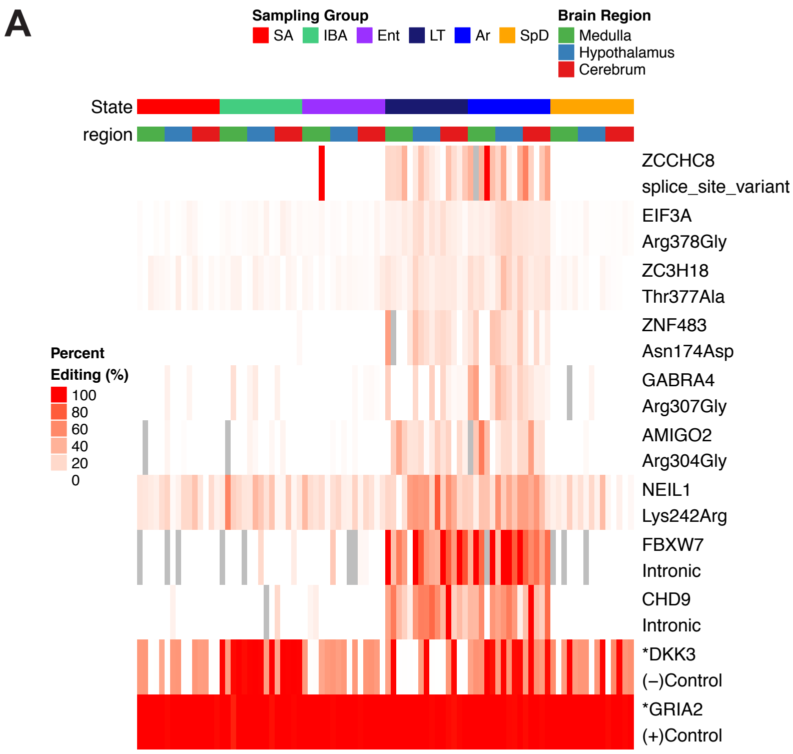

#### Supplemental Figure 8

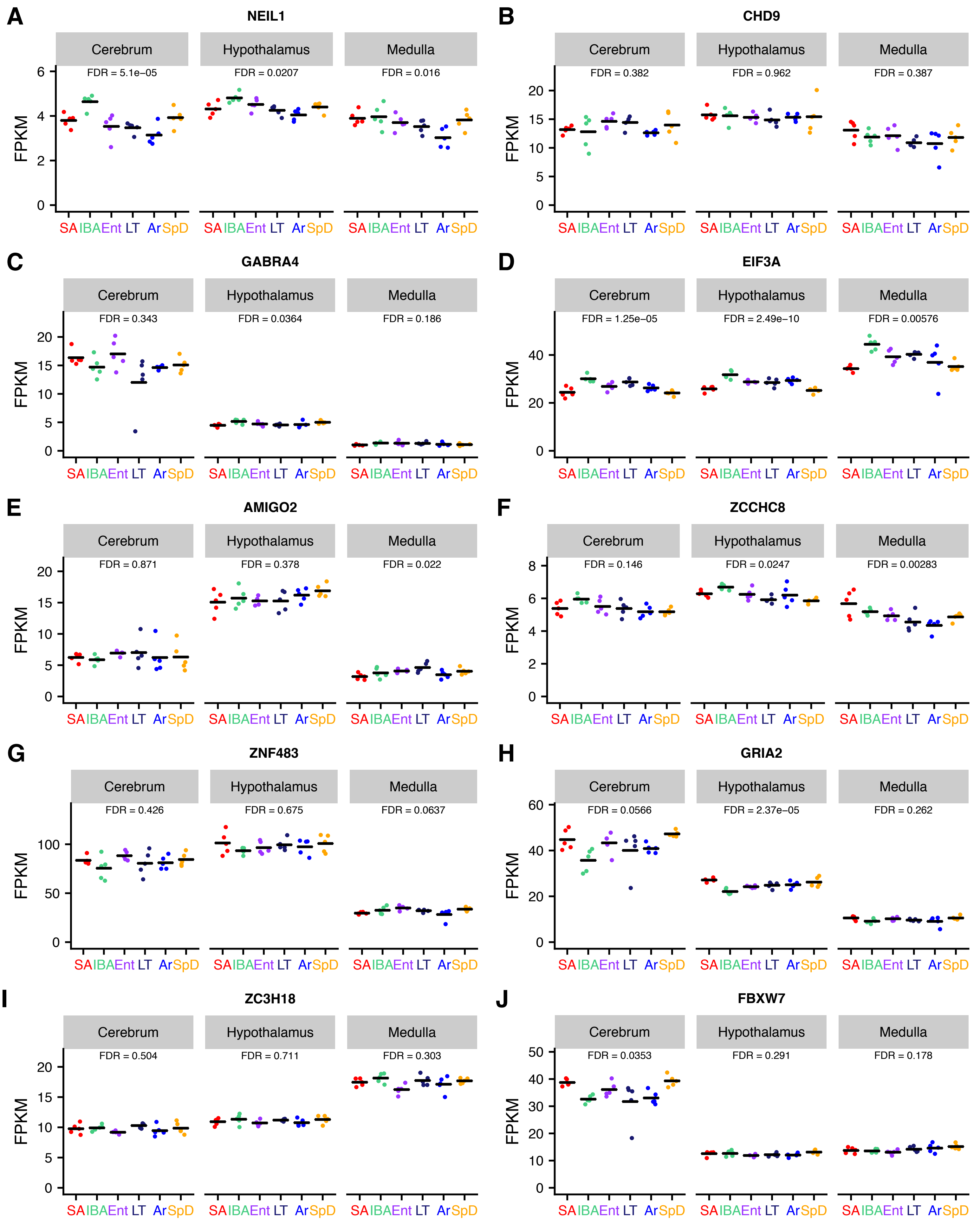

#### Supplemental Figure 9

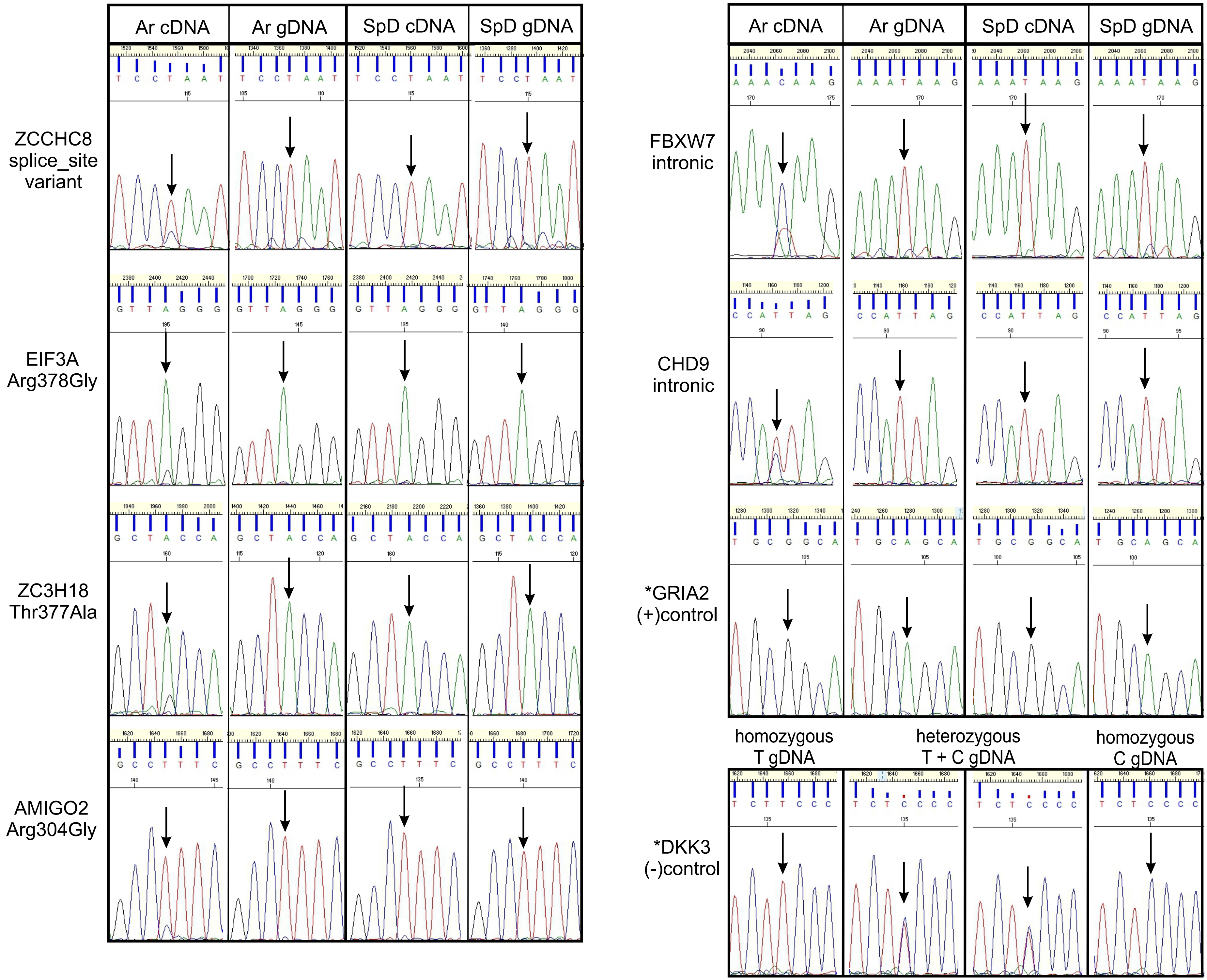

#### Supplemental Figure 10

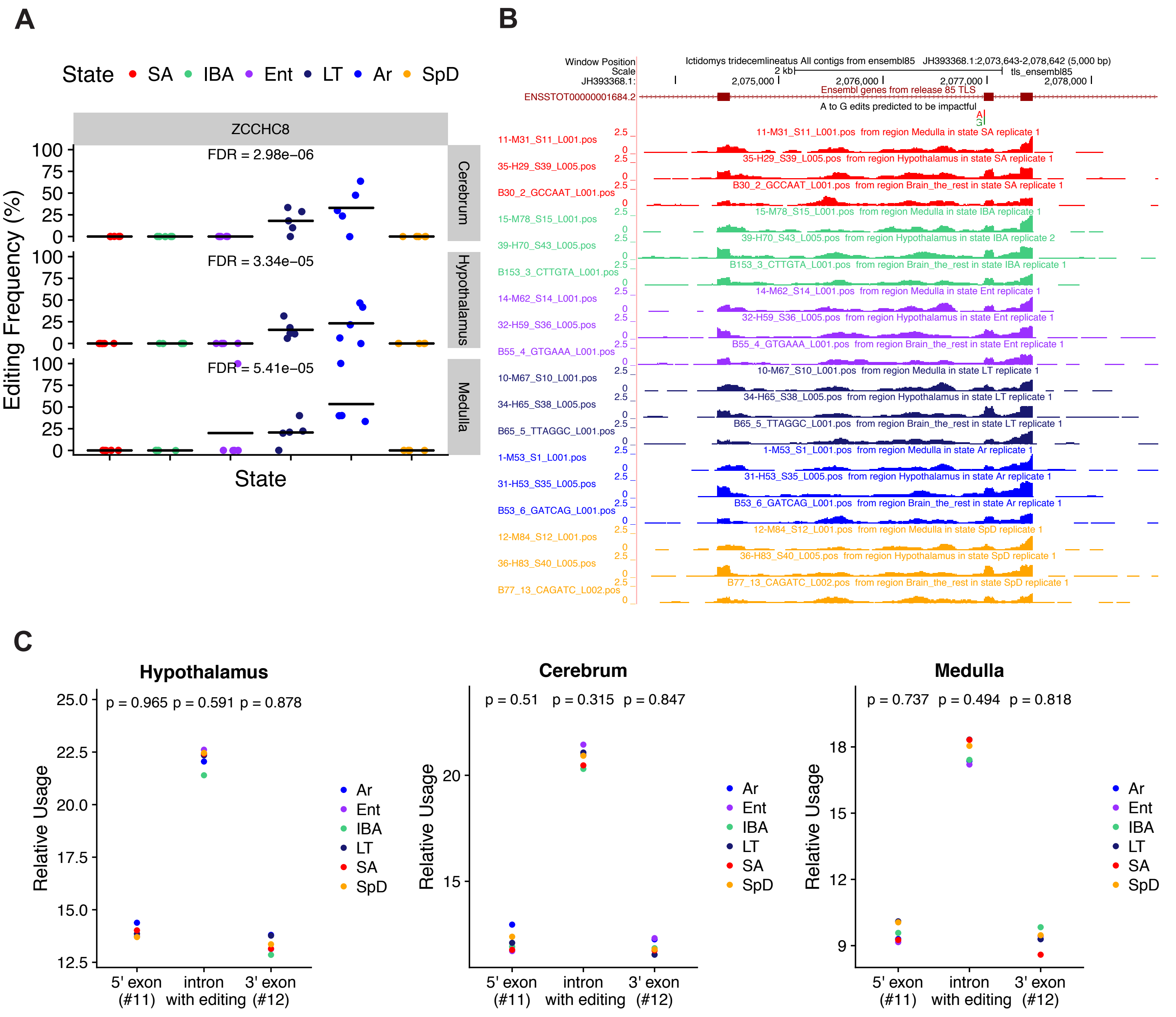

**Supplemental Figure 11**

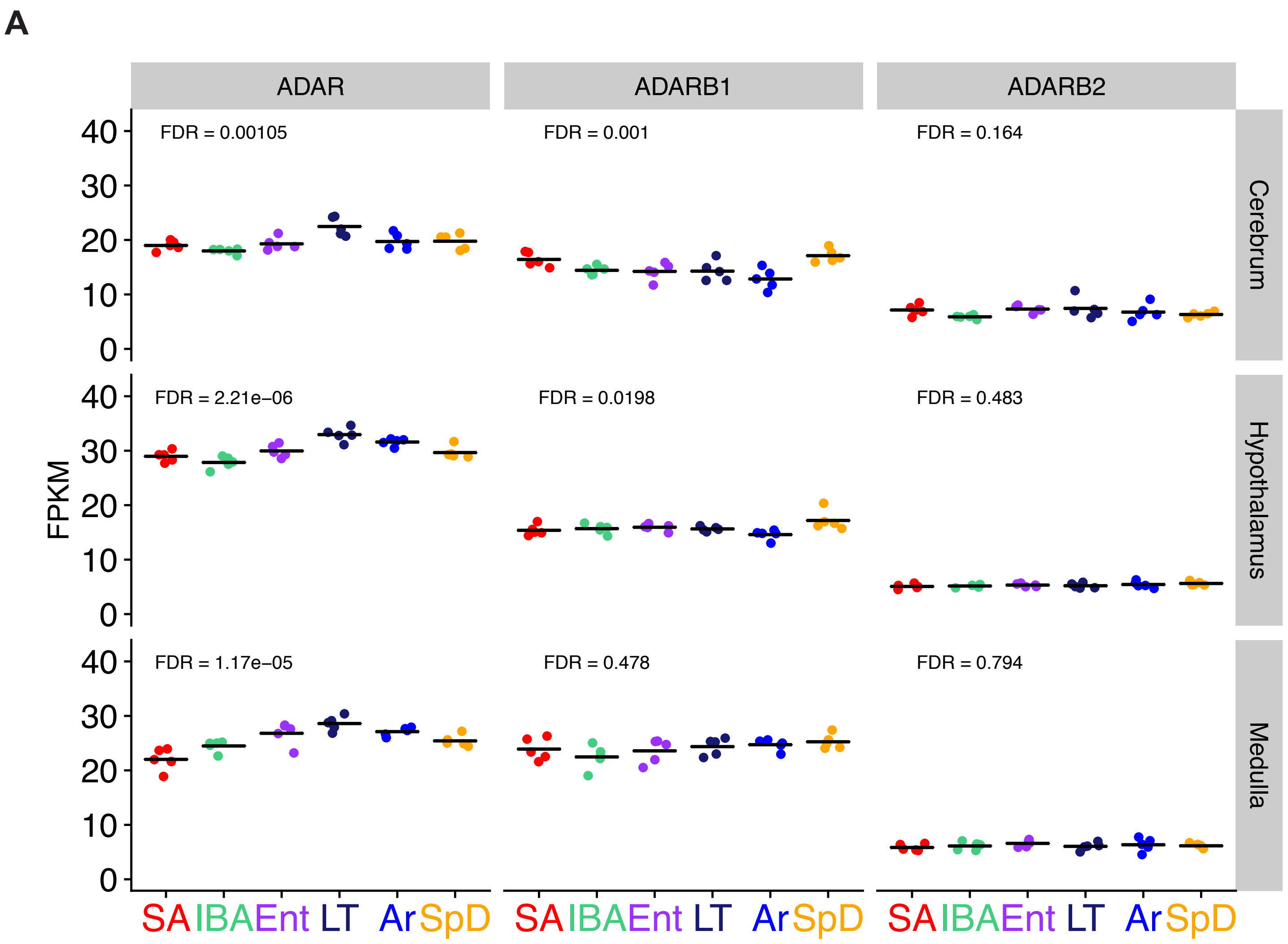

**Supplemental Figure 12**

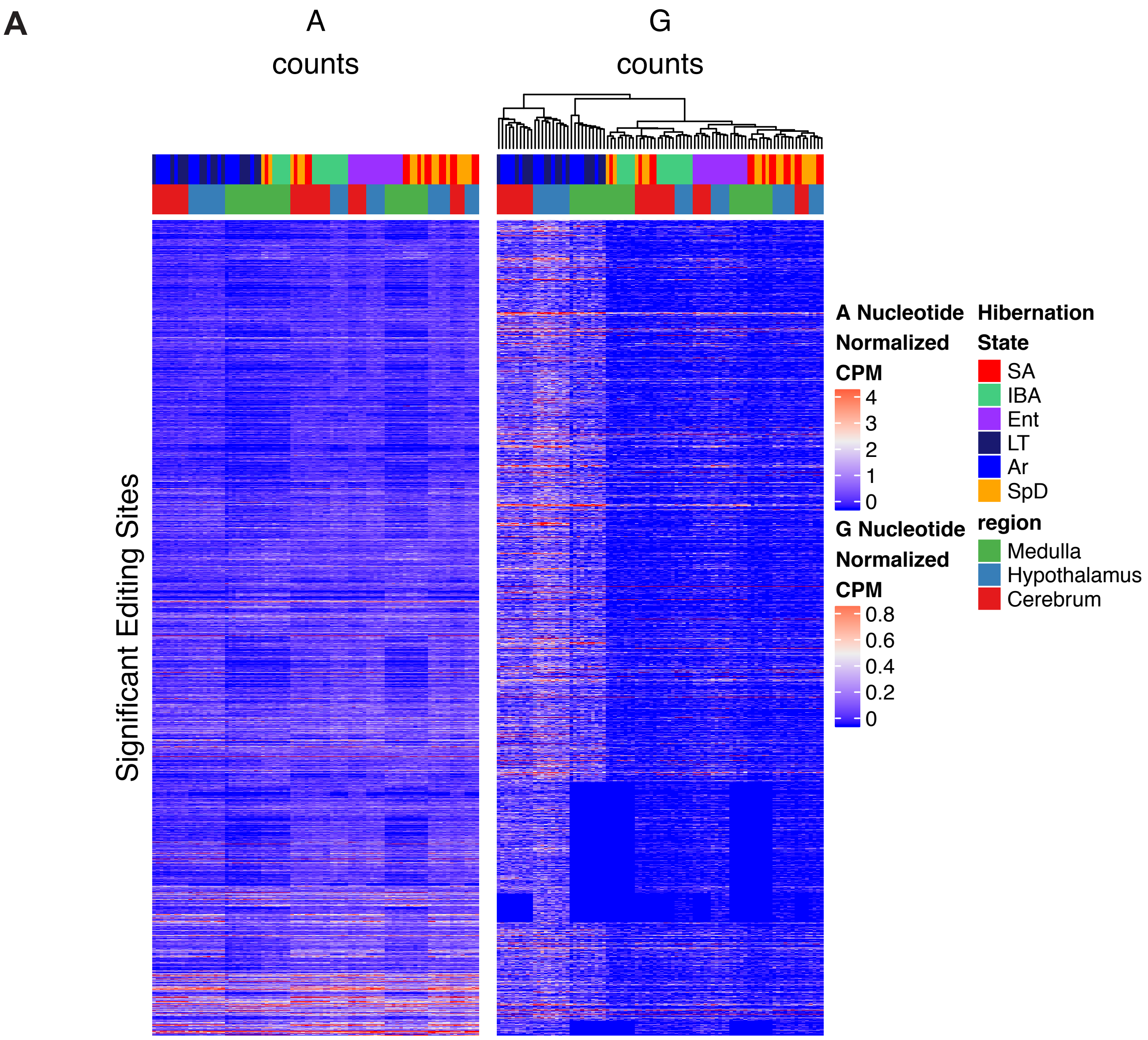
